# Parallel adaptive responses to postponed reproduction increase lifespan and immune defense

**DOI:** 10.64898/2026.02.25.708047

**Authors:** Karen Alma Gamboa-Santarosa, Giovanni A. Crestani, Alejandro Moran, Dolly Modha, Hannah S. Dugo, Mansour Abdoli, Molly K. Burke, Parvin Shahrestani

## Abstract

Experimental evolution studies with *Drosophila melanogaster* have long played a role in the effort to dissect the genetic basis of aging and longevity. While selection for postponed reproduction reliably extends lifespan, additional phenotypic consequences and the genomic bases of this adaptation remain unclear. Here, we leveraged the highly replicated *Drosophila* Experimental Evolution Population (DEEP) system to further investigate the relationship between longevity and other life-history traits. Derived from the same wild population, two tenfold-replicated treatments, each with populations derived from two different ancestral backgrounds: a control treatment in a 14-day generation cycle was maintained for 56 generations, and an experimental treatment in a 70-day cycle maintained for 20 generations. Experimental populations evolved to have longer lifespan, delayed development time, increased fecundity, greater stress resistance, and stronger immune defense. Pooled-population genomic data reveal highly convergent allele frequency shifts within treatments, and point to 300 candidate genes underlying differentiated phenotypes. Candidate genes are enriched for functional categories involving neural development and morphogenesis rather than canonical aging or immune defense pathways. These results recapitulate that selection for postponed reproduction drives broad physiological changes and a highly polygenic adaptive response, with an unprecedented level of experimental replication.

## INTRODUCTION

Natural populations harbor substantial genetic variation for lifespan and aging, defined as the post-reproductive decline in survival and fertility. Evolutionary theory provides a theoretical framework for understanding why lifespan is limited and variable: the force of natural selection declines with age (Hamilton, 1966), so alleles with late-acting deleterious effects may accumulate in populations (Medawar, 1952), while alleles with early-life benefits can be favored despite later costs (Williams, 1957). These processes generate segregating genetic variation for lifespan, and predict that longevity can evolve. Indeed, classic selection experiments with *Drosophila melanogaster* and other species demonstrate that selection for postponed reproduction consistently produces longer-lived populations (reviewed by McHugh and Burke, 2022; Rose et al., 2004), and this postponed longevity is accompanied by correlated shifts in other life-history traits.

Postponed longevity has been repeatedly evolved in *Drosophila* across laboratories (e.g. Luckinbill et al., 1984; Partridge and Fowler, 1992; Remolina et al., 2012; Rose, 1984; Zwaan et al., 1995). Within tens of generations, evolved populations exhibit altered reproductive schedules and improved stress-resistance phenotypes in addition to longer lifespans (reviewed by Burke and Rose, 2009). More recently, Evolve and Resequence, or E&R (Long et al., 2015; Turner et al., 2011) strategies have been applied to characterize the genetic basis of these traits. E&R studies consistently show that the genetic architecture of aging-related traits is highly polygenic and involves subtle frequency shifts at many loci (Burke et al., 2010; Carnes et al., 2015; Fabian et al., 2018; Graves et al., 2017; Hoedjes et al., 2019). However, candidate genes differ across studies, with little overlap and few canonical longevity genes implicated by any of them (explored by Fabian et al., 2018; Hoedjes et al., 2019). This inconsistency likely reflects variation in founder populations, experimental design, and selection regimes, but could also stem from low statistical power.

Most E&R longevity studies employ modest replication (3-5 populations), small population sizes (N_e_<1000), and relatively short experimental durations. Simulations and empirical work suggest that replication is the critical parameter for linking phenotypes to genomic loci (Baldwin-Brown et al., 2014; Burke, Liti, et al., 2014; Kofler & Schlötterer, 2014). Underpowered designs obscure subtle but replicable genomic responses, limiting the ability to distinguish true targets of selection from background noise. Thus, the lack of overlap across studies could be systematic of the high false positive and false negative rate associated with insufficient replication, or it could implicate genuine biological contingency. Highly replicated designs provide a powerful route for evaluating the repeatability of evolution and for testing whether selection for postponed reproduction consistently targets particular physiological processes.

One process of growing interest in the context of experimentally evolved longevity is immune defense. While most E&R longevity studies have not implicated canonical aging genes, Fabian et al., 2018, reported differentiation at loci associated with immune defense between long-lived and control populations. This observation raises the possibility that immune defense, or the response after the exposure to a pathogen, may be an underappreciated axis of adaptation to postponed reproduction. Conceptually, this link is plausible. Immune activation can confer substantial early-life benefits with pathogenic challenge, yet chronic or excessive immune signalling can accelerate senescence, a pattern broadly consistent with antagonistic pleiotropy (Schwenke et al., 2016, Franceschi et al., 2000). Across taxonomic groups, mounting or maintaining immune responses is energetically costly, and investment in immunity is often coupled with shifts in reproductive allocation, stress physiology, and somatic maintenance. These life-history trade-offs position immune defense as both a potential early-life enhancer of fitness and also a late-life liability, aligning neatly with the evolutionary theory of aging. With this in mind, investigators have begun to examine the relationship between immune defense and longevity in *Drosophila* through the lens of experimental evolution, though the extent to which each trait evolves as a correlated response to selection for the other therefore remains unclear. When immune defense is the focal trait under direct selection, enhanced immunity appears to come at the cost of reproductive output and lifespan (Shahrestani et al., 2021). Alternatively, when age of reproduction is the focal trait under direct selection, improvements to longevity and reproduction lead to correlated improvements in immune defense without apparent costs (Bagheri et al., 2025), and differentiated loci implicate genes related to immune function (Burke et al. 2014; Fabian et al. 2018). These mixed outcomes underscore the complexity of interactions between immunity and aging, and highlight the need for more powerful, highly-replicated tests.

Here, we leveraged the *Drosophila* Experimental Evolution Population (DEEP) system (Bagheri et al., 2025) to conduct an E&R experiment with unprecedented replication. The DEEP system descends from Rose’s (1984) classic founding populations and includes tenfold replicated treatments of postponed reproduction (“O-type”) versus baseline controls (“B-type”). Our goals were to: i) characterize the phenotypic consequences of selection for postponed reproduction, with an emphasis on immune defense; and ii) describe the genomic architecture of this adaptation by comparing allele frequencies across the first 20 generations of O-type selection. We predicted a polygenic basis, with convergence across replicate populations and ancestral backgrounds (Graves et al., 2017). We observed longer lifespan, increased fecundity and body weight, longer development times, greater stress resistance, and notably stronger immune defense. Genomic analyses revealed highly polygenic, convergent adaptation, implicating genes not linked to canonical aging pathways or immune defense, but to neural development and morphogenesis.

## METHODS

### DEEP Experimental Evolution System

All populations in the *Drosophila* Experimental Evolution Population (DEEP) System were generated from wild *D. melanogaster* sampled from an apple orchard in South Amherst, MA (the “IV”/Ives population) that have been maintained in the laboratory under controlled conditions for nearly 50 years (Ives, 1970). This ancestral IV population is maintained on discrete 14-day generation cycles and was used to establish five “baseline” populations (B_1-5_) and five long-lived populations (O_1-5_) via experimental evolution for postponed reproduction (Rose, 1984). The B_1-5_ populations are maintained on 14-day generation cycles, while in the O_1-5_ populations, the age of first reproduction was progressively postponed until generation cycles could be routinely kept at 70 days. In 2008, a new set of populations was created from the O_1-5_ treatment; their regime was reverted to the ancestral 14-day generation cycles, and they were named BO_1-5_ (Burke et al., 2016). The B_1-5_ and BO_1-5_ populations rapidly converged phenotypically (Burke et al., 2016) and at the genomic level (Graves et al. 2017).

In October 2018, B_1-5_ and BO_1-5_ populations were transferred from Dr. Michael Rose’s laboratory at the University of California, Irvine (UCI), to Dr. Parvin Shahrestani’s laboratory at the California State University, Fullerton (CSUF). At CSUF, the B_1-5_ and BO_1-5_ populations are maintained under nearly-identical conditions to those at UCI, including a banana-molasses diet, 24-hour light cycle, constant 25°C temperature, cage enclosures, and large census population sizes (>1,000). All 20 populations are reared in vials for 12 days before we dump each vial in its own cage. Six generations after this transition, we derived two treatments called nB_1-5_ and OB_1-5_ from each B_1-5_ population, and two treatments called nBO_1-5_ and OBO_1-5_ from each BO_1-5_ population (Figure 1). The control treatments nBO_1-5_ and nB_1-5_ remained as “B-type”(on 14-day generation cycles), with the “n” designator standing for “new” and being added to indicate that these populations are maintained outside the ancestral UCI environment. In these populations, flies were transferred from vials into cages on day 12 from egg, given yeasted food plates on day 13, and had their eggs collected from them on day 14. On the other hand, the “O” designator signals experimental populations selected for delayed reproduction, culminating in a 70-day generation cycle. They are referred to as the “O-type” populations. The transition from B-type to O-type was done gradually to minimize the chance of a population crash (i.e. population sizes were kept at >1,000 individuals each generation), and to maximize the genetic diversity preserved. This gradual increase involved two generations of 21-day cycles, two generations of 28-days cycles, five generations of 35-day cycles, two generations of 42-day cycles, two generations of 56-day cycles, two generations of 63-day cycles, and finally 70-day cycles for each subsequent generation.

**Figure 1.**
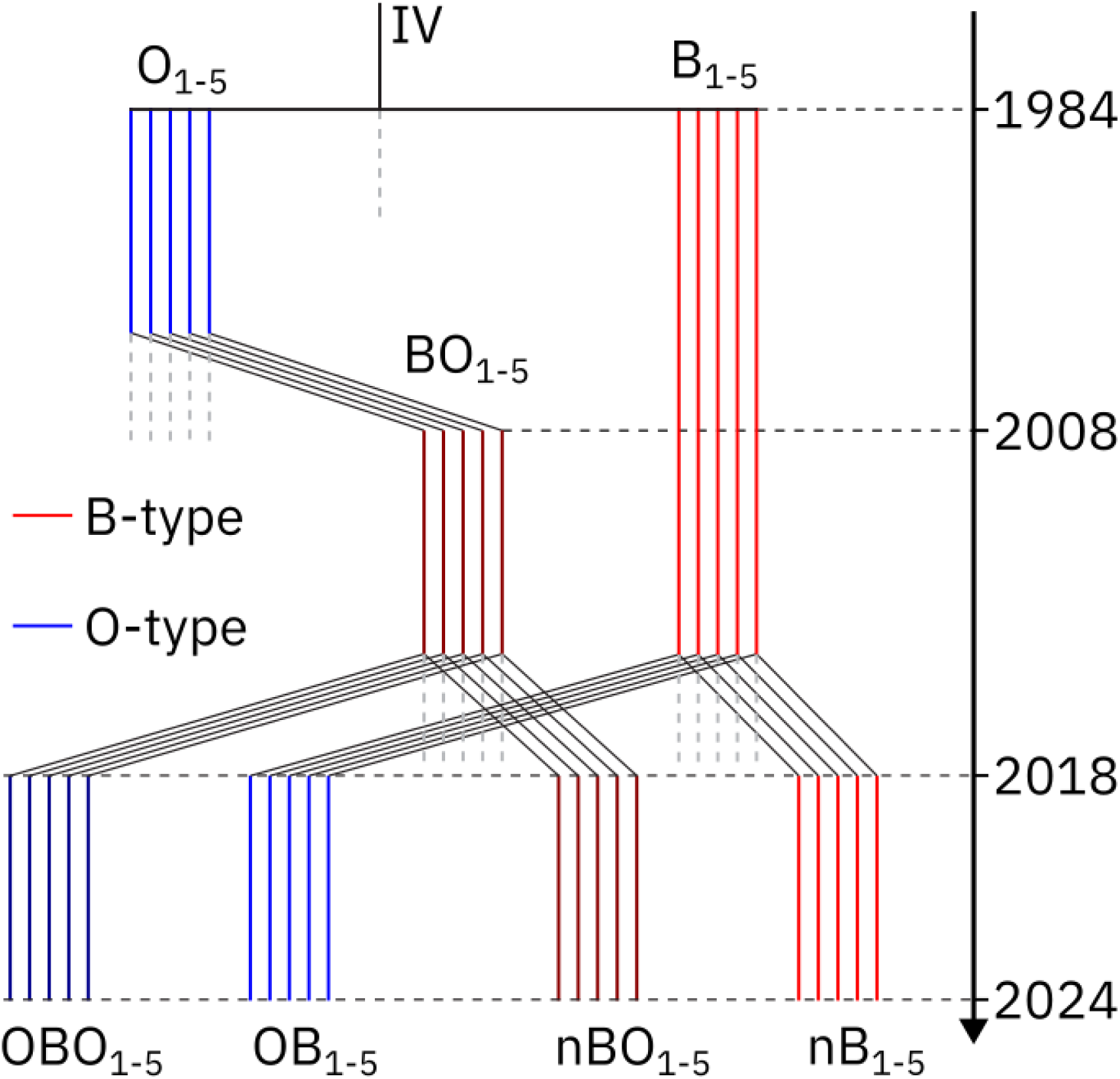
Schematic of the evolutionary history of experimental populations under O-type and B-type selection. Populations represented by blue lines experienced O-type selection whereas populations represented by red lines experienced B-type selection. Dashed gray lines indicate periods of population continuity not relevant to the current study. The splitting of populations (e.g. O_1-5_ into BO_1-5_, and B_1-5_ & BO_1-5_ into the final 20 populations) happened instantaneously and therefore violates the y-axis scale. **ALT TEXT:** Phylogeny of the twenty populations used in the study, from 1984 to 2024, with color showing selection regimen.

The 20 populations (nBO_1-5_, OBO_1-5_, nB_1-5_, and OB_1-5_) constitute a valuable resource for exploring *Drosophila* life-history traits. With five-fold replication for each ancestry within each regime (B-derived vs. BO-derived), and ten-fold replication for each longevity regime (B-type vs. O-type), this system offers a rich set of possible comparisons. Figure 1 summarizes these populations, their ancestry, and their current maintenance regime.

### Phenotyping

We surveyed life-history, stress resistance, and immune phenotypes in all 20 experimental populations after approximately 20 generations of O-type selection. Longevity, development time, fecundity, and body weight were assayed at exactly generation 20, and immune defense and starvation/desiccation resistance were assayed at generation 22. For all phenotype assays, flies were sampled from each population and cycled through two 14-day generations prior to the assay to minimize parental or environmental effects. Adult flies were assayed at day 14 from egg except where otherwise indicated. We also performed, with identical methods, a longevity and a development time assay at an intermediate timepoint (generation 12 of O-type selection), and data are presented at Supplementary Figure 1. We summarize each assay procedure below. Additional details and expanded protocols can be found in Walsh (2022).

#### Age-specific mortality rate & longevity

To quantify age-specific mortality and lifespan, we established three replicate cages for each experimental population, each initiated with around 500 flies (60 cages total). Dead flies were sexed, counted, and removed from cages four times a week until all flies died in all cages. For a given day *d*, mortality rate was calculated with the following equation:

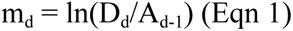

where D_d_ is the number of flies that died on that day and A_d-1_ is the number of flies alive at the end of the previous day. Mean population longevity was calculated by averaging mean cage longevity across the three cages representing each experimental population.

#### Development time

Following a pre-laying period, adults from each population were provided fresh yeasted food plates (one per cage) and allowed three hours to lay eggs. Batches of 60-80 eggs were transferred from each food plate into each of five vials of fresh banana medium (100 total vials across all populations). Vials were monitored throughout their pupal development, and adult eclosion was recorded beginning from the emergence of the first fly from its pupal case in each vial. Vials were checked every 6 hours until all eggs eclosed. Mean development time per vial was calculated and then averaged across all five vials to determine mean population-level development time.

#### Fecundity

For each population, we established 10 vials with a removable cap containing yeasted charcoal medium (to facilitate egg counting), into which we transferred four male and four female flies. Flies were anesthetized daily to replace the charcoal medium, and the number of eggs on each day-old charcoal plug was recorded at the same time daily. Vials retained four male and four female flies throughout the experiment. Dead flies were sexed and replaced (using CO2 anesthesia) with a fly of the same sex sourced from a designated “Replacement” (R) replicate vial, which would always be the highest numbered replicate vial (e.g. nB_1_-vial10 would be relabeled nB_1_-vial10R after being used to refill another vial). Once an R-vial was depleted, the next highest-numbered replicate (e.g. CB_1_-vial9) became the new source. Egg counts from R-vials were included in the analysis only for the day before their first use as a source; subsequent egg counts were excluded. All R-vials were maintained under the same daily handling and feeding conditions as experimental vials.

#### Body weight

Body weight was measured for flies aged 17 days from egg. For each population, 50 females and 50 males were sorted into groups of 10 same-sex individuals and weighed in pre-weighed microcentrifuge tubes. Mean vial weight was calculated for each sex and then averaged across replicate vials to obtain mean population weight. Additional measurements of dry weight and water loss from related assays are reported in Walsh (2022).

#### Starvation and Desiccation Resistance

To assay starvation resistance, we collected 20 females and 20 males at age 15 days from egg. Groups of five same-sex individuals were anesthetized and transferred to vials (80 vials total), which were capped with a cotton ball soaked with 5 mL of DI water beneath a foam-plastic plug cut in half. We sealed vials with two layers of parafilm to conserve internal humidity. This setup denied flies access to food, while providing enough moisture to prevent death from dehydration. Vials were monitored every four hours and deaths recorded as they occurred. An identical protocol was used for desiccation resistance assays, replacing wet cotton balls with 3 g of desiccant per group. Vials were monitored every hour and deaths recorded as they occurred. For both assays, we calculated mean population resistance by averaging mean vial resistance across all replicate vials of a given population.

#### Immune Defense

To quantify immune defense we exposed flies to the fungal entomopathogen *Beauveria bassiana* strain GHA (Bioworks, Inc., Victor NY, lot number TGA1-96-06B). Flies were sampled randomly from rearing vials and divided into two replicate infected and two replicate control cages, with 100 to 200 flies per cage (estimated by volume during sampling), for a total of 80 cages. Infected cages were sprayed with a solution of 0.3 g of fungal spores suspended in 25 mL of a 0.03% Silwet surfactant solution, which was shaken by hand and a wrist-action shaker for 15 minutes. The control cages were sprayed with a solution of DI water and surfactant. For each spray, anesthetized flies were spread out on Petri dish lids and placed on ice to keep them immobile. 5mL of the appropriate spray was applied using a custom-built spray tower (Shahrestani et al., 2021) designed to introduce controlled, even doses to the fly cuticle. Sprayed flies were transferred to population cages and maintained at 100% humidity for 24 hours to promote fungal germination and cuticle penetration. The humidity was then lowered to ∼60% for the remainder of the assay, and fly mortality was monitored daily for 12 consecutive days. Infected and control death percentages were calculated by averaging across the two replicate cages per population.

#### Phenotypic Statistical Analysis

All statistical analyses were conducted in R version 4.5.0 (R Core Team, 2025). Data manipulation used the packages readxl v1.4.5 (Wickham & Bryan, 2015) and tidyverse v2.0 (Wickham et al., 2019), and ggplot2 v3.5.2 (Wickham, 2016) was used for visualization. Survival analyses were performed using survival v3.8-3 (Therneau et al., 2024) and survminer v0.5.1 (Kassambara et al., 2025). For mortality rate, we used Cox Proportional Hazard regression (Cox, 1972) via survival::coxph, modeling survival time as a function of “Regime”, “Sex”, and “Ancestry”. Proportionality assumptions were tested using survival::cox.zph, and all parameters failed the test. Hence, the model was adjusted to allow the effect (hazard ratio) of the coefficients to change over experimental time. Differences in population mean values for other phenotypes were analyzed using linear models or generalized linear models when normality assumptions were violated. Fixed effects included “Regime”, “Sex”, and “Ancestry”, with “Infection” added for immune defense. We considered the interaction terms “Infected:Regime” when analyzing Immune defense data, and “Ancestry:Regime” when analyzing all phenotypes. Welch’s t-tests for population mean comparisons (Figure 2 and Supplementary Figure 2) were conducted using the package ggpubr v0.6.1 (Kassambara, 2016). Full statistical analysis output tables are available as Supplementary Table 1.

**Figure 2.**
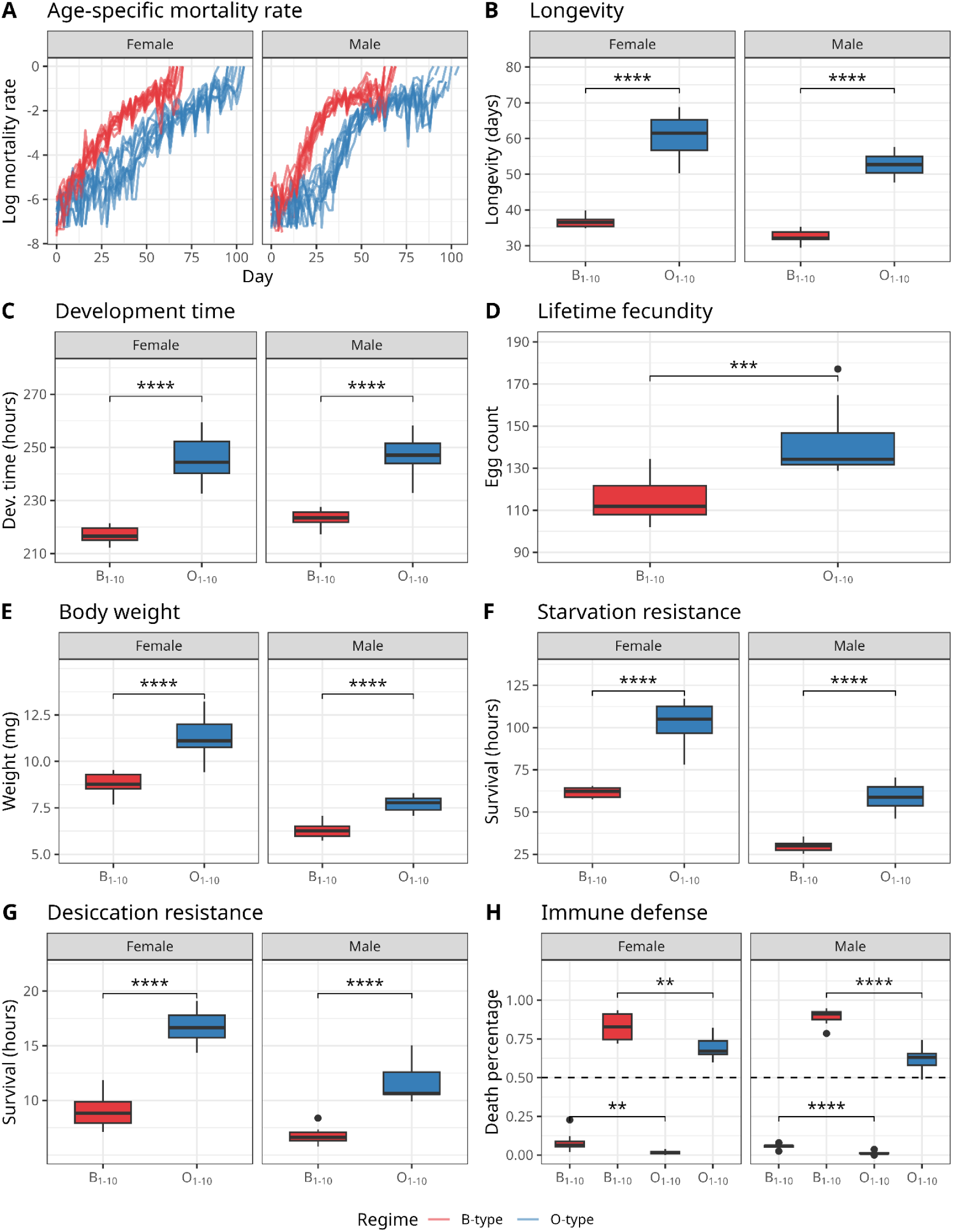
B-type and O-type selection regimes lead to differentiation in life-history and stress-resistance phenotypes. Across all panels, phenotypes measured in the B-type populations (nBO_1-5_ combined with nB_1-5_) are shown in red, and phenotypes measured in the O-type populations (OBO_1-5_ combined with OB_1-5_) are shown in blue. Each boxplot in panels B-H is derived from 10 data points (n=10 for all groups), with each data point representing a single value per replicate population per sex. Female and male data are shown separately where applicable. Statistical significance of the effect of selection regime was assessed using t-tests on population mean values, comparing B-type and O-type regimes. **A** Age-specific mortality rate over time; each line represents one replicate population. **B** Mean longevity (days). **C** Mean development time (hours). **D** Mean lifetime fecundity (average number of eggs per female). **E** Mean bodyweight (mg). **F** Mean starvation resistance (survival time in hours). **G** Mean desiccation resistance (survival time in hours). **H** Immune defense, measured as percent mortality following infection (death percentage); plots above the dashed line show infected files, while plots below the dashed line show uninfected control flies. **ALT TEXT:** Graphs comparing phenotype of O-type flies and B-type flies, with statistical significance markers.

### Sequencing and Bioinformatic Processing

#### Library preparation

Live flies were sampled from each of the 20 populations on day 14 from egg at generations 01 and 20 of O-type selection, and cryopreserved in ethanol at -80°C. Genomic DNA from 100 female flies per replicate per treatment per timepoint (40 pools total) was extracted using Qiagen (Puregene) reagents, following the vendor’s protocol for bulk extraction. We prepared sequencing libraries using a high-throughput tagmentation protocol (modified Illumina Nextera library preparation protocol), pooling the 40 uniquely barcoded libraries into a single multiplex. This multiplexed library was run on three lanes of a NextSeq2000 (P2, PE150) at the Oregon State University Center for Quantitative Life Sciences. Low-coverage samples were identified, re-pooled (from the same original group of 100 females), and re-sequenced on an additional two lanes to obtain higher and more consistent coverage across all populations.

#### Genomic Pipeline

We modified an existing GATK-based Pool-SEQ pipeline (Phillips et al., 2020) to identify SNPs and small INDELs relative to the *D. melanogaster* genome (v. 6.51), using Nextflow (Di Tommaso et al., 2017) to increase efficiency and reproducibility. We aligned reads using BWA MEM v0.7.18 (Vasimuddin et al., 2019) with the default program parameters and called variants with GATK v4.2.0.0 (Auwera & O’Connor, 2020), following best practices (Poplin et al., 2018). Then, we filtered the resulting VCF file with GATK VariantFiltration using the following filtering parameters: “QD < 5.0, MQ < 40.0, FS > 60.0, ReadPosRankSum < -8.0, MQRankSum < -12.5”.

We converted this merged VCF file into a SNP table containing AD (allele depth) and DP (unfiltered depth) as the alternate allele count and coverage for all SNPs passing the quality filters. To ensure high-quality data for downstream analysis, we filtered out SNPs with coverage below 50 or above 500 in at least one sample (Schlötterer et al., 2014). Additionally, we imposed a minor allele frequency filter to exclude SNPs with a combined frequency of 1% across all sampled populations, eliminating both variants fixed in all samples (uninformative) and spurious variants likely caused by sequencing or variant calling errors.

#### Allele frequency analysis

After quality filtering, we calculated allele frequencies by dividing the alternative allele count (AD) by the coverage (DP). We then performed a Principal Component Analysis (PCA) (function stats::prcomp) in R to visualize the extent to which ancestry versus selection treatment shaped genome-wide allele frequencies. We applied a K-means clustering algorithm (k = 3) on allele frequency data to verify grouping patterns. We chose k=3 because we expected three groups, one with experimental samples at generation 20, and one per ancestry containing experimental samples at generation 01 and control samples. To identify alleles associated with the observed phenotypes, we applied an adapted CMH test (Spitzer et al., 2020) on a scaled SNP table (we set DP to 100 for all SNPs, and adjusted AD proportionally). Scaling is applied to control for coverage heterogeneity. We corrected for multiple testing bias (FDR-corrected), and considered the SNPs with adjusted p-values below a threshold of 1^-100^ as statistically significant (roughly top 0.1% quantile).

#### SNP annotation and Gene Ontology (GO) term analysis

We used TxDb.Dmelanogaster.UCSC.dm6.ensGene v3.12.0 (Bioconductor Core Team, 2017), AnnotationDbi v1.70.0 (Hervé Pagès, 2017), and GenomicRanges v1.60.0 (Lawrence et al., 2013) to to obtain a list of genomic features containing significant SNPs. To explore the GO terms associated with the genes, we used the R packages org.Dm.eg.db v3.21.0 (Carlson, 2017) and biomaRt v2.64.0 (Durinck et al., 2009). We excluded GO terms annotated with the TAS, NAS, IC, and ND codes from the analysis, as these are considered low confidence annotation codes. To identify enriched Gene Ontology (GO) terms at the Biological Process level, we used the package clusterProfiler v4.16.0 (G. Yu, 2024), more specifically the function clusterProfiler::enrichGO set to a q-value cutoff of 0.05 and to only capture “Biological Process” terms. To reduce redundancy, we used the clusterProfiler::simplify function with default parameters. We used the package enrichplot v1.28.4 (Guangchuang Yu, 2018) to plot the enriched GO terms. Our background gene set was all *Drosophila melanogaster* genes, which was provided by the package.

## RESULTS

### Phenotypes

Across all phenotypic assays, O-type flies consistently differed in trait values relative to the B-type controls after 20 generations (and after 10 generations, as shown in Supplementary Figure 1). They displayed longer lifespans, delayed development, higher fecundity, and increased resistance to starvation, desiccation, and fungal infection. These patterns indicate a broad phenotypic shift associated with selection to extended generation times. All analyses were conducted at the population level (N = 20 populations per model).

#### Mortality rate & longevity

O-type flies had significantly lower age-specific mortality rates (Figure 2A, Supplementary Figure 2A) than B-type flies in both sexes (Cox Proportional-Hazards test, P<0.001). Male flies had a higher hazard rate of death (Cox Proportional-Hazards test, P<0.001). Consistent with this pattern, mean population longevity (Figure 2B, Supplementary Figure 2B) was estimated to be higher in O-type flies when compared to B-type flies (linear model, *P*<0.001), and lower in male flies when compared to female flies (linear model, *P*<0.001). Data for an intermediate time point are available as Supplementary Figure 1.

#### Development time

Population development time (Figure 2C, Supplementary Figure 2C) was significantly longer in O-type flies compared to B-type flies (linear model, *P*<0.001). However, this difference was smaller in O-type flies with BO ancestry (linear model interaction term, *P*=0.005). Intermediate generation data are available as Supplementary Figure 1.

#### Fecundity

O-type populations exhibited higher mean population fecundity (Figure 2D, Supplementary Figure 2D) at all ages compared to B-type populations (Poisson generalized linear model, *P*<0.001). Also, we observed a weak significant effect of ancestry for the fecundity phenotype, such that mean population fecundity in the BO-derived populations was estimated to be higher than in the B-derived populations, regardless of selection regime (Poisson generalized linear model, *P*<0.058). Age-specific fecundity trajectories are displayed in Supplementary Figure 3.

#### Body weight

O-type flies had higher estimated mean population weight than B-type flies (linear model, *P*<0.001, Figure 2E, Supplementary Figure 2E), and female flies were estimated to have higher mean population weight than male flies (linear model, *P*<0.001). Dry weight and water loss data are reported in Walsh, 2022.

#### Starvation Resistance and Desiccation Resistance

O-type flies showed increased resistance to starvation and desiccation, with a higher proportion of flies surviving under both stresses compared to B-type flies. O-type flies had higher mean population starvation resistance (Figure 2F, Supplementary Figure 2F) than B-type flies (linear model, *P*<0.001). Also, females were estimated to have higher mean population starvation resistance than males under starvation conditions (linear model, *P*<0.001). For desiccation resistance (Figure 2G, Supplementary Figure 2G), O-type flies were estimated to have higher mean population desiccation resistance than B-type flies (linear model, *P*<0.001) and females were estimated to have higher mean population desiccation resistance than males (linear model, *P*<0.001).

#### Immune defense

O-type flies showed a higher probability of surviving *Beauvaria bassania* infection compared with B-type flies (Figure 2H, Supplementary Figure 2H). O-type flies were estimated to have a reduced chance of dying if compared to B-type populations, regardless of infection status or sex (Binomial generalized linear model, *P*<0.001). Infection predictably reduced survival, with Infected flies having higher odds of dying when compared to non-infected Control flies (Binomial generalized linear model, *P*<0.001), and O-type selection regime being estimated to have increased survival in the infected flies (Binomial generalized linear model interaction term, *P*=0.038), indicating that O-type flies experienced less infection-induced mortality than B-type flies.

### Genomics

After filtering steps, the resulting dataset consisted of 1,086,149 biallelic SNPs across the five major chromosome arms. The mean sequencing genome-wide coverage ranged from 56X to 183X, with a mean of 106X across all samples. Coverage was reasonably consistent across the genome, with a coefficient of variation ranging from 0.20 to 0.30.

PCA analysis showed the landscape of allele frequencies across all populations (Figure 3). We observed patterns of convergence based on selection treatment: O-type populations clustered together regardless of ancestry, based on similar patterns of genetic variation after 20 generations. Populations at generation 1 clustered according to ancestry.

**Figure 3.**
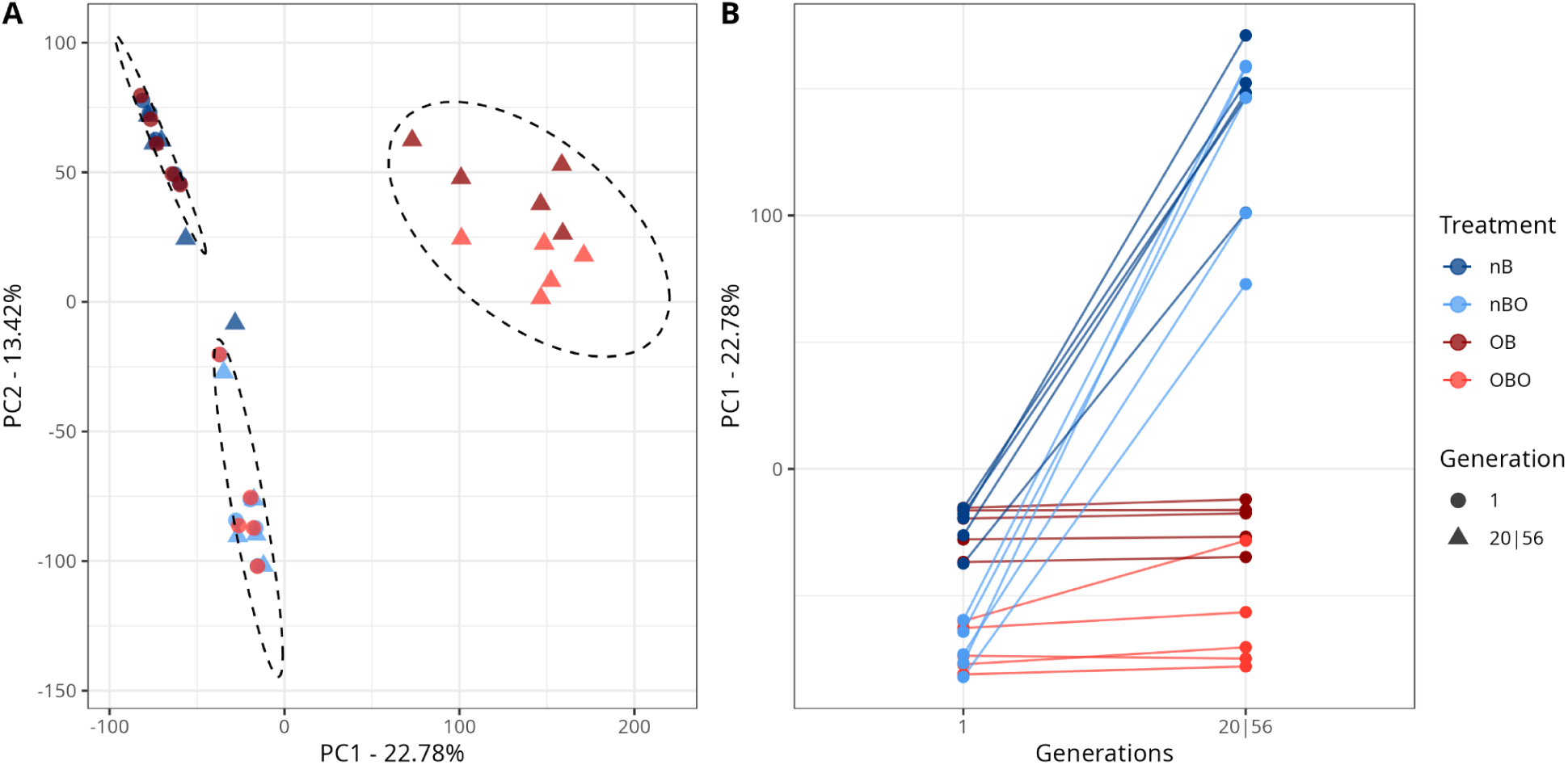
Genome-wide allele frequencies converge by selection regime rather than ancestry after 20 generations of selection. Principal component analysis (PCA) of SNP allele frequencies (1,086,149 filtered SNPs) across all populations shows that evolved populations cluster by selection regime rather than ancestry. **A** K-means clustering (k=3) of allele frequencies. The top left cluster contains OBO_1-5_ populations at generation 1 and nBO_1-5_ populations at generations 1 and 56, grouping all non-evolved populations derived from BO_1-5_. The bottom left cluster contains OB_1-5_ at generation 1 and nB_1-5_ at generations 1 and 56, grouping all non-evolved populations derived from B_1-5_. The right cluster contains OBO_1-5_ at generation 20 and OB_1-5_ at generation 20, grouping all evolved experimental populations. **B** Principal component 1 plotted by generation. By generation 20, experimental populations diverge from their ancestral and control populations, which remain clustered together, consistent with convergence under shared selection regimes. **ALT TEXT:** Scatterplot of first two principal components, with samples grouping by treatment and generation, and line plot showing principal component one by generations.

As the PCA analysis indicated convergence between OBO_1-5_ and OB_1-5_, we performed CMH tests comparing the allele frequencies between generations 01 and 20 of all the O-type populations combined (referring to them as O_1-10_). Benjamini-Hochberg FDR corrected, log-transformed p-values were plotted in Figure 4A. We identified a few dozen defined peaks evenly distributed across all 4 main chromosome arms and X, with 1,218 statistically significant SNPs, located within 290 unique genes (240 of which have documented annotations). The annotated list of candidate SNPs is available in Supplementary Table 2, and the list of candidate genes is available in Supplementary Table 3. When analyzed individually, CMH test results considering only the OBO and OB populations largely implicated the same regions, but with highly reduced statistical significance (Figure 4, panels B and C). When comparing B_1-10_ between generations 01 and 55, we observe three peaks that are located in unannotated genomic regions (Supplementary Figure 4). These peaks are not shared with the other comparisons. When comparing O_1-10_ at generation 20 with B_1-10_ at generation 56, we observe differentiation in similar regions as when comparing O_1-10_ across generations, albeit with less statistical support (Supplementary Figure 5).

**Figure 4.**
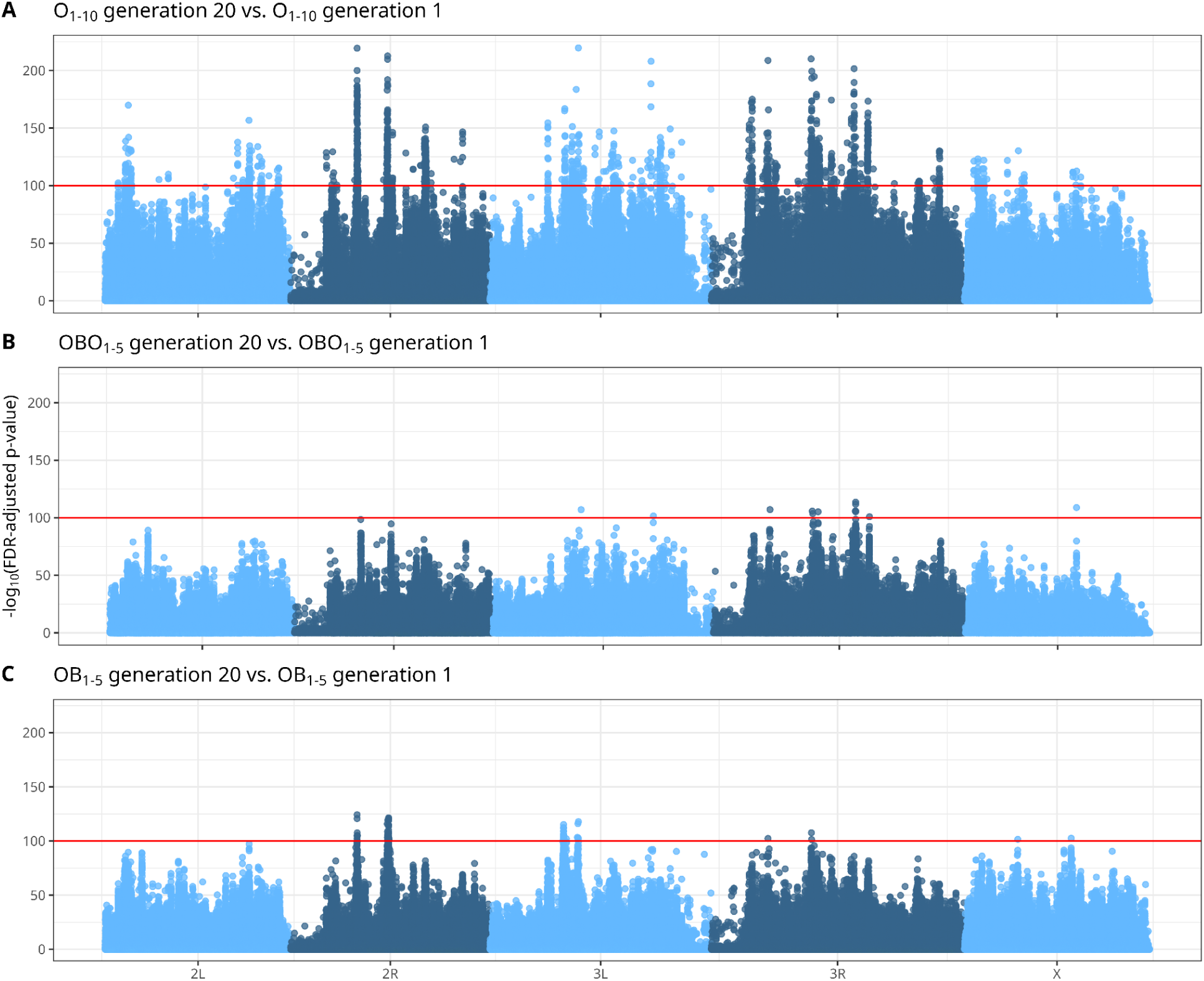
Combining replicate O-type populations increases power to detect evolved genome-wide allele frequency change. Manhattan plots show Cochran-Mantel-Haenszel (CMH) test results based on scaled SNP frequencies; x-axis values represent cumulative physical distance along the genome and y-axis values represent FDR-corrected *p*-values. The horizontal red line indicates the significance threshold (adjusted *p*-value = 10^-100^). **A** Combined analysis of the ten O-type populations (OBO_1-5_ & OB_1-5_, relabeled here as O_1-10_) at generation 20 compared to generation 1. **B** OBO_1-5_ populations at generation 20 compared to generation 1. **C** OB_1-5_ populations at generation 20 compared to generation 1. **ALT TEXT:** Manhattan plots of O-type populations compared between generations 1 and 20, with significance threshold indicating statistically significant single nucleotide polymorphisms.

Out of the 290 candidate genes, 240 genes have annotated GO terms, 214 of which are Biological Process terms. A GO term enrichment analysis suggested an enrichment for 42 terms (Supplementary Table 4) over the 290 unique genes implicated by our candidate SNP list (Supplementary Table 2). Removing redundant terms (see Methods for more details) returns a consolidated list of 21 terms (Figure 5). The terms GO:0048667 (cell morphogenesis involved in neuron differentiation), GO:0048812 (neuron projection morphogenesis), GO:0120039 (plasma membrane bounded cell projection morphogenesis), GO:0048858 (cell projection morphogenesis), and GO:0031175 (neuron projection development) had the lowest q-values (< 0.0001) and were enriched in 26 of the 214 considered genes. Figure 5 shows the simplified enriched GO terms with adjusted p-values and ratios. Though we observed no GO term enrichment for any longevity-related term, six genes on the list have been previously associated with the “determination of adult lifespan” (GO:0008340) GO term (or any child term): *Adcy1* or *rut* (Tong et al., 2007), *Egfr* (Kamakura, 2011), *mthl5* (Araújo et al., 2013), *Gnmt* (Obata & Miura, 2015), *Lkb1* (Funakoshi et al., 2011), and *puc* (Libert et al., 2008; M. C. Wang et al., 2003). Similarly, we observe 9 genes previously associated with the “Immune response” (GO:0006955) (or any child) GO term: *snk* (Irving et al., 2001), *dia* (Howell et al., 2012), *Atf3* (Rynes et al., 2012), *Spn88Ea* (Ahmad et al., 2009), *ths* (Dragojlovic-Munther & Martinez-Agosto, 2013), *Gbp1* (Tsuzuki et al., 2012), *tefu* (Petersen et al., 2012), *Elp6* (Howell et al., 2012), *puc* (Libert et al., 2008), *Antp* (Mandal et al., 2007), and *sick* (Foley & O’Farrell, 2004).

**Figure 5.**
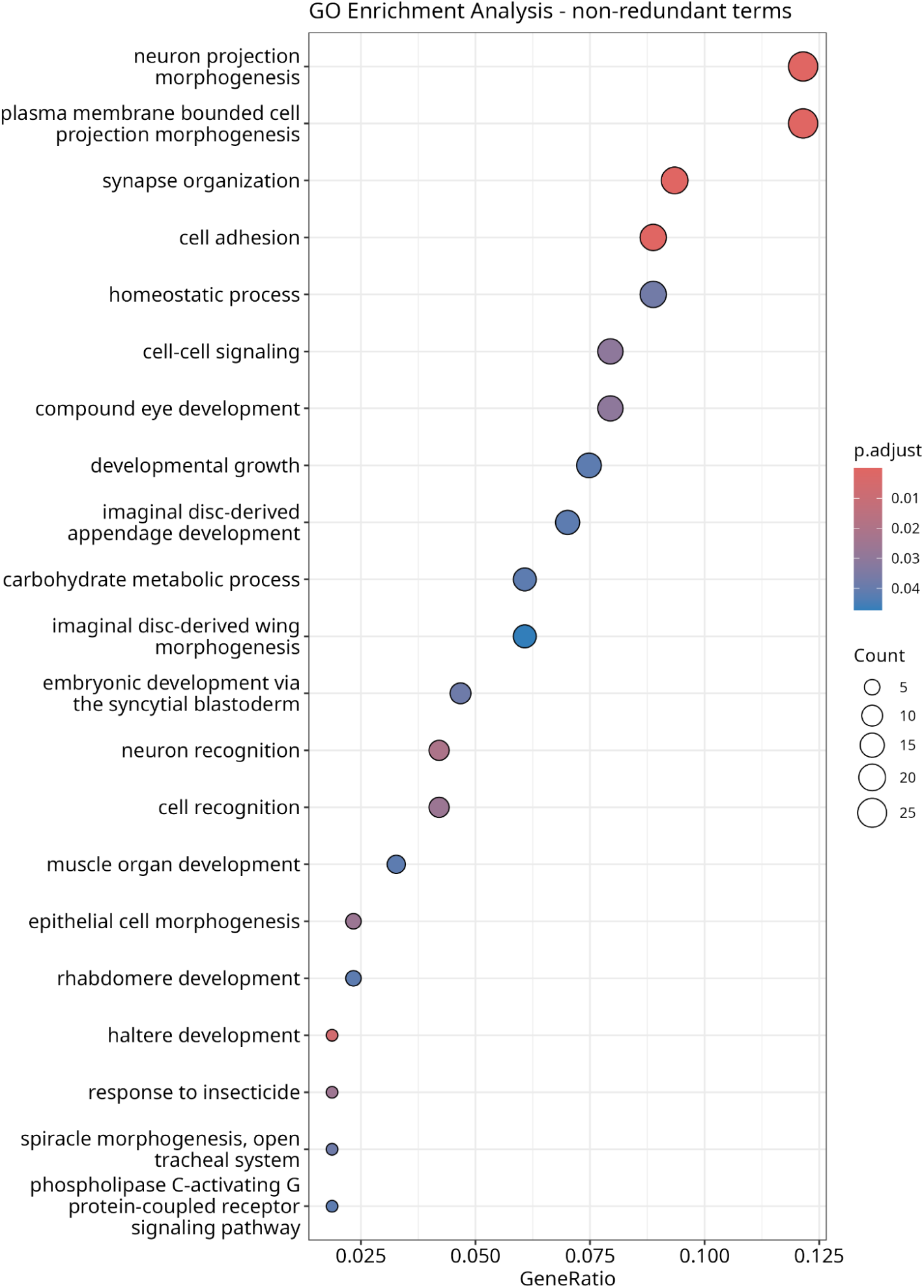
Enriched biological process GO terms among genes containing significant SNPs. The y-axis lists enriched Biological Process GO terms. Point size corresponds to the number of genes associated with each term, and point color indicates the Benjamini-Hochberg FDR adjusted *p*-value correction). The x-axis shows the gene ratio, defined as the number of genes associated with a given GO term divided by the total number of genes analyzed. **ALT TEXT:** Graph showing gene enriched ontology terms, with datapoint color depicting P-value and datapoint size depicting gene count.

## DISCUSSION

### Phenotypic consequences of selection for postponed reproduction

Under O-type selection, all populations consistently evolved extended longevity along with several correlated traits: increased development time, fecundity, immune defense, body weight, and resistance to starvation and desiccation. These phenotypic changes align with expectations from prior experimental evolution work (*cf.* Rose et al., 2004) and confirm that postponed reproduction imposes strong selection on life-history traits in the DEEP system.

Notably, O-type flies exhibited higher fecundity at all ages compared to B-type flies (Supplementary Figure 3). This contrasts with earlier findings of a trade-off between longevity and early-life reproduction (Rose, 1984). This discrepancy may reflect experimental differences, such as dietary yeast supplementation, which has been shown to mitigate early fecundity costs (Chippindale et al., 1993). O-type flies also survived longer than B-type flies under starvation and desiccation stress. As body weight influences the response to these stressors (Djawdan et al., 1998), the observed increased weight in O-type flies may underlie their enhanced resistance. This increase in weight may have resulted from prolonged larval feeding (associated with the increase in development time in O-type populations). However, since we only measured time to eclosion, we cannot determine at which developmental stage the B- and O-type flies diverge. Follow-up experiments dissecting the developmental timeline in finer detail could clarify whether differences stem from the larval or pupal period.

Overall, phenotypic evolution under O-type selection led to expected increases in organismal robustness, with no detectable trade-offs aside from increased development time, which could incur ecological costs under natural conditions.

### Absence of trade-offs between immune defense and longevity

Trade-offs between fecundity and immune defense are well-documented in insects (Schwenke et al., 2016), suggesting that a physiological investment in reproduction can weaken immune defense, while immune activation reduces fecundity. Relating immune defense to longevity, Shahrestani et al. (2021) reported that *Drosophila* populations selected for improved defense against *Beauveria bassiana* exhibited higher post-infection survival but reduced lifespan under uninfected conditions. Thus, reduced immune defense in O-type populations would be consistent with a trade-off hypothesis; yet, we observed consistently improved measures of immune defense across all O-type populations. Our observations instead align with studies showing positive associations between longevity and immune performance. Fabian et al. (2018) reported that populations with experimentally-evolved extended lifespan also exhibited improved immune defense against multiple pathogens, including *B. bassiana*. Bagheri et al. (2025) similarly found no evidence of trade-offs between longevity and immune defense in other DEEP populations. Together, these observations contribute to the ongoing narrative that correlated traits don’t always respond similarly when one or the other is selected upon directly (reviewed by Burke and Rose 2009), and they also suggest that correlations between immune defense and longevity depend on the nature of the immune strategies favored by selection.

Recent conceptual work emphasizes that immune defense is not a single trait, but rather a suite of strategies that differ in their physiological costs and evolutionary consequences (e.g. Troha and Ayres 2022). Antagonistic immune strategies, such as chronic or constitutive immune activation, can impose substantial costs through immunopathology or energetic expenditure, whereas cooperative or condition-dependent immune strategies scale with organismal condition and may enhance resistance or tolerance with fewer trade-offs. Selection for postponed reproduction may favor the latter, particularly if extended lifespan increases cumulative exposure to pathogens in crowded cage environments.

In addition, O-type populations show increased adult body weight, presumably due to increased larval feeding over their extended development time, which may further contribute to enhanced immune defense. Mounting an immune response is an energetically costly process that remodels metabolism to inhibit nutrient storage and catalyze fat stores (DiAngelo et al., 2009). In *Drosophila*, the fat body functions as a central immune and metabolic organ, integrating nutrient storage with systemic immune signaling (reviewed by Yu et al. 2022). Because fat body mass and lipid reserves scale with overall body condition, variation in body size or energetic state should influence immune performance. Consistent with this framework, larger and more robust O-type flies may show enhanced immune performance because immune defense scales with overall condition rather than being constrained by zero-sum allocation trade-offs. However, because immune defense was measured at a single age (14 days from egg), the relationship between body condition, immune defense, and longevity across the lifespan remains unresolved. Future experiments assaying immune defense across multiple ages, multiple pathogens or infection intensities will be necessary to more comprehensively examine the impact of evolved longevity on age-specific immune competence.

### Effects of ancestry on phenotypic evolution

Although selection consistently drove robust phenotypic divergence between O-type and B-type populations, we detected a few ancestry-associated differences when comparing BO-derived (OBO_1-5_) and B-derived (OB_1-5_) lines within treatments (Supplementary Figure 2). This finding generally aligns with previous observations that populations derived from this experimental system rapidly converged at both phenotypic (Burke et al., 2016) and genomic levels (Graves et al., 2017), suggesting that strong directional selection can quickly overwhelm consequences of evolutionary history. Our statistical models only weakly support ancestry as being a significant factor in a limited set of comparisons. The presence of ancestry effects across multiple life-history traits suggests that convergence under experimental evolution regimes is not uniformly rapid, but instead may proceed at different rates depending on the trait and the evolutionary context. Among these traits, fecundity stands out as the most consistently affected and therefore provides a useful point of comparison to prior work.

Burke et al. (2016) found that early-life fecundity did not fully converge between selection treatments that differed in long-term, but not recent evolutionary history. Specifically, early fecundity of “ACO” populations did not fully converge with “AO” populations after the latter had transitioned from a 70-day to a 9-day generation cycle over ∼160 generations. Given that our O-type populations have experienced substantially fewer generations of selection, the persistence of ancestry-associated fecundity differences, and by extension, differences in correlated traits, is consistent with a lag in convergence rather than a failure of parallel evolution. In contrast, Burke et al. (2016) reported full convergence of early fecundity between B-type treatments that differed in their long-term, but not recent evolutionary history (the same B and BO lines used to initiate the nB and nBO lines here). The observation of fecundity differences between nB and nBO populations in the present study is therefore less intuitive. One potential explanation is that Burke et al. (2016) quantified early fecundity (ages 11-14 days from egg), whereas we measured lifetime fecundity. It is possible that lifetime reproductive output, or age-specific fecundity later in life, did not fully converge in the earlier experiment, and that effect has continued to persist. Additionally, in the present study, nB populations were transitioned to cage environments prior to the onset of the experiment, whereas nBO populations had long been maintained in cages. Such environmental differences could plausibly influence reproductive schedules, potentially reintroducing detectable phenotypic differences even among populations that had previously converged under earlier assay conditions.

Taken together, these results do not contradict the broader conclusion of Burke et al. (2016) that strong selection rapidly drives phenotypic convergence. Rather, they indicate that convergence may be trait-specific and/or age-specific, and likely shaped by the strength of selection imposed by the reproductive regime. Under the more extreme O-type conditions, which impose strong selection by restricting reproduction to late life, populations have rapidly diverged from the B-type phenotype but may require additional generations to fully converge with one another. In contrast, under the more permissive B-type conditions, weaker selection may allow residual ancestry effects or context-dependent differences to persist. These patterns are also reflected in the genomic comparisons among populations as described below.

### Genomic consequences of selection

Experimental evolution for postponed reproduction produced widespread and consistent genomic changes in outbred *Drosophila* populations, consistent with a polygenic response. Despite clear phenotypic divergence in longevity and immune defense, our genomic analysis did not reveal overrepresentation of genes traditionally associated with those life-history traits. Instead, the strongest signals emerging from allele frequency change over 20 generations of O-type selection were found in genes associated with GO-terms related to development, and particularly neural development (Figure 5).

Our list of candidate genes does not include many canonical “aging loci” identified through mutant screens, such as *mth* (Petrosyan et al., 2014), *Indy* (P.-Y. Wang et al., 2009), *chico* (Slack et al., 2015), *mTor* (Kapahi et al., 2004), etc. However, we found seven genes previously associated with “determination of adult lifespan”, and nine genes previously associated with “immune response”. Among these, only one (growth-blocking peptide *Gbp1*) is involved with antimicrobial peptides (AMPs), a group of molecules critical in the innate immune response. Unlike studies based on inbred lines and *de novo* mutations, our experimental system applies long-term selection to genetically diverse populations, allowing polygenic adaptation via subtle shifts in allele frequency across many loci. In this context, our observed enrichment in Biological Process terms suggests that selection may have favored genes involved in maintaining physiological homeostasis and somatic maintenance, consistent with generalized robustness phenotypes rather than targeted aging pathways.

Comparing our gene list to those of other investigators working with long-lived lines experimentally evolved for increased longevity reveals relatively little overlap among our studies. Carnes et al. (2015) identified 99 candidate genes in females, eight 8.08%) of which match our candidate genes (*Amy-d*, *Gem3*, *ko*, *Ccdc85*, *CG11400*, *Gbp2*, *CG45263*, and *asRNA:cr45272*). Given that this study used populations derived from the same founding Ives population, we might have expected even more overlap. *Gbp2* is a notable gene on this list; the GBP pathway is suggested as an important pathway in mediating acute innate immune reactions (Tsuzuki et al., 2012), and *Gbp2* regulates *Gbp1* expression (Ono et al., 2024). In a study with populations that come from a completely independent study system, Fabian et al. (2018) identified 868 candidate genes, with twenty (2.18%) also being present in our list (*Ace*, *Atf3*, *CG13280*, *CG1677*, *CG33203*, *CG33459*, *CG34354*, *CG42673*, *comm3*, *DIP-gamma*, *kirre*, *Lar*, *MICU3*, *myd*, *Nepl19*, *path*, *sqz*, *ths*, *Ugt37D1*, and *wat*). No genes are simultaneously reported as candidates by the three studies, highlighting the challenge of drawing connections across similar experiments at the gene level. Differences in the methods used to assess significance and define candidate gene lists likely contributed to this limited overlap.

### Role of replication and ancestry in genomic interpretation

One strength of our design is 10-fold replication among O-type populations, which enables us to detect consistent selection responses despite the modest effect sizes typical of polygenic traits. Previous studies with fewer replicates may have lacked the power to identify these subtle, repeatable patterns, which may help explain why candidate genes often differ across experiments.

Compared to what we observed among phenotypes, ancestry played an even subtler role in shaping the genomic response. Comparisons between OBO_1–5_ and OB_1–5_ revealed minimal divergence, reinforcing the idea that strong selection can drive parallel genomic changes across distinct ancestral backgrounds. This convergence at the genomic level suggests that key targets of selection are shared across genetic backgrounds, despite variable phenotypic expression.

We also compared genomic signals across the B-type populations through generations (Supplementary Figure 4). While we observed negligible differentiation between the nB_1-5_ populations over time, three weak peaks emerged from comparing the nBO_1-5_ populations initially and after 55 generations. These three peaks persist when grouping all 10 B-type populations together and comparing them over time, though they become non-significant with the increased replication. While these three noncoding regions could implicate effects of domestication and ongoing selection in the nBO lines, they may also reflect differences in mapping quality. This further emphasizes the importance of biological replication in the genomic interpretation of selection experiments.

We also compared signals of genomic differentiation between O_1-10_ at generation 20 with B_1-10_ at generation 56 (Supplementary Figure 5). This analysis implicated broadly similar genomic regions to those identified in comparisons among O_1-10_ across generations, though with reduced statistical power. The weaker signal is consistent with the expectation that control populations are themselves evolving under laboratory conditions, including responses to domestication and husbandry that are independent of the O-type life-history regime. As a result, contrasts that rely exclusively on contemporary control populations may incorporate genomic change unrelated to the focal selective pressure, potentially obscuring loci most relevant to adaptation to O-type selection. This interpretation is consistent with prior results demonstrating substantial evolutionary change in laboratory control populations (e.g. Phillips et al. 2016) and highlights the value of longitudinal sampling in Evolve & Resequence experimental design.

### Conclusions and future directions

Our study demonstrates that selection for postponed reproduction consistently drives both phenotypic and genomic changes in *D. melanogaster*, with broad convergence across replicate lines. Diverged phenotypes largely recapitulate those observed in prior evolution experiments, though our method of assaying immune defense is novel in this study system. Our observation that immune defense is enhanced in O-type populations is therefore noteworthy and invites deeper lines of inquiry. Links between longevity and innate immunity are well-established, and the DEEP population resource provides a tantalizing opportunity to explore the connections between these two complex traits. While our genomic results do not directly implicate genes in known *Drosophila* immune pathways, future work, particularly the integration of gene expression and other -omics data types, should enrich our understanding of these traits in outbred populations.

## Data availability

The data underlying this article are available in the Dryad Digital Repository, at https://doi.org/10.5061/dryad.mcvdnckf5. Data analysis R scripts are available at https://github.com/gacrestani/Gamboa-Santarosa_et_al. Raw sequencing data is available at SRA PRJNA1392952.

## Author contributions

**Karen Alma Gamboa-Santarosa:** Formal analysis, Investigation, Validation, Writing – original draft.

**Giovanni A. Crestani:** Formal analysis, Investigation, Validation, Visualization, Writing – original draft.

**Alejandro Moran:** Investigation.

**Dolly Modha**: Investigation.

**Hannah S. Dugo:** Investigation.

**Mansour Abdoli:** Formal analysis.

**Molly K. Burke:** Conceptualization, Data curation, Funding acquisition, Methodology, Project administration, Resources, Supervision, Validation, Writing – review & editing.

**Parvin Shahrestani:** Conceptualization, Data curation, Funding acquisition, Methodology, Project administration, Resources, Supevision, Validation, Writing – review & editing.

## Acknowledgements

We thank Dr. Michael Rose (UC Irvine) for generously providing the ancestral populations that were used to found this experiment. We also thank Avani Modha from Dr. Parvin Shahrestani’s (CSUF) laboratory for her notable contributions as well as the entire Longevity Selection Team. We declare the usage of ChatGPT (models GPT-4 and GPT-3.5) and Google Gemini (models 2.0 Flash and 3.0 Pro) for initial stages of data analysis and figure generation to streamline R coding (subsequently reviewed and verified before usage) and sentence-level editing of the manuscript’s text.

## SUPPLEMENTARY INFORMATION

**Supplementary Figure 1.**
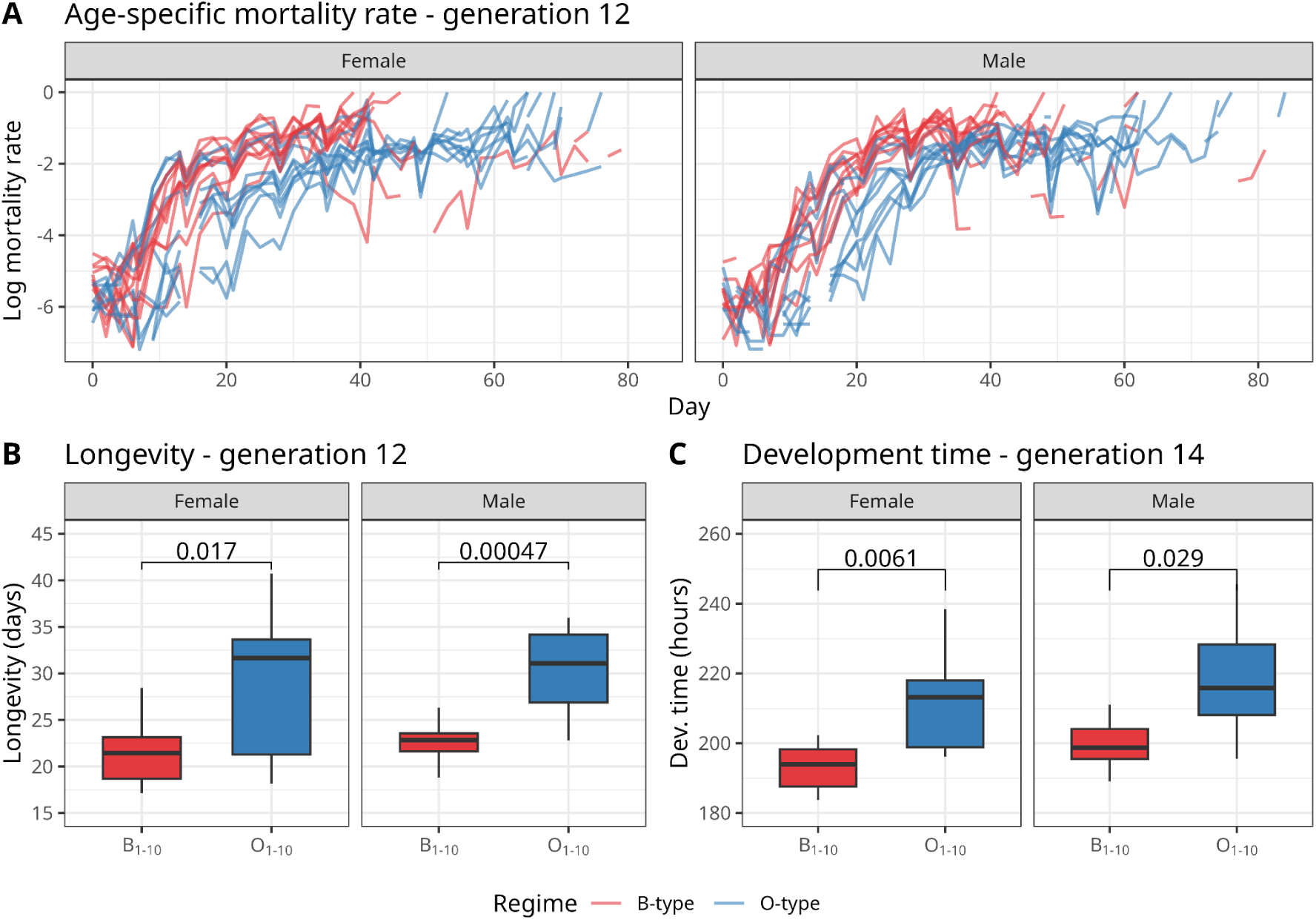
Phenotypic divergence is evident after ∼12 generations of O-type selection. Phenotypic data were collected at an intermediate timepoint, corresponding to generations 12 and 14 of O-type selection and approximately 19 generations of B-type selection. Across all panels, data from the B-type regime (nBO_1-5_ combined with nB_1-5_) are shown in red, and data from the O-type regime (OBO_1-5_ combined with OB_1-5_) are shown in blue. Panels B and C, show boxplots derived from 10 data points (n=10 for all groups), with each data point representing a sex-specific population mean from each replicate population. Data are shown separately for males and females. **A** Instantaneous mortality rate over time; each line represents one replicate population, with plots split by sex. **B** Mean population longevity (days). **C** Mean development time (hours). **ALT TEXT:** Graphs of phenotypes measured at an intermediate generation time point, with statistical comparisons.

**Supplementary Figure 2.**
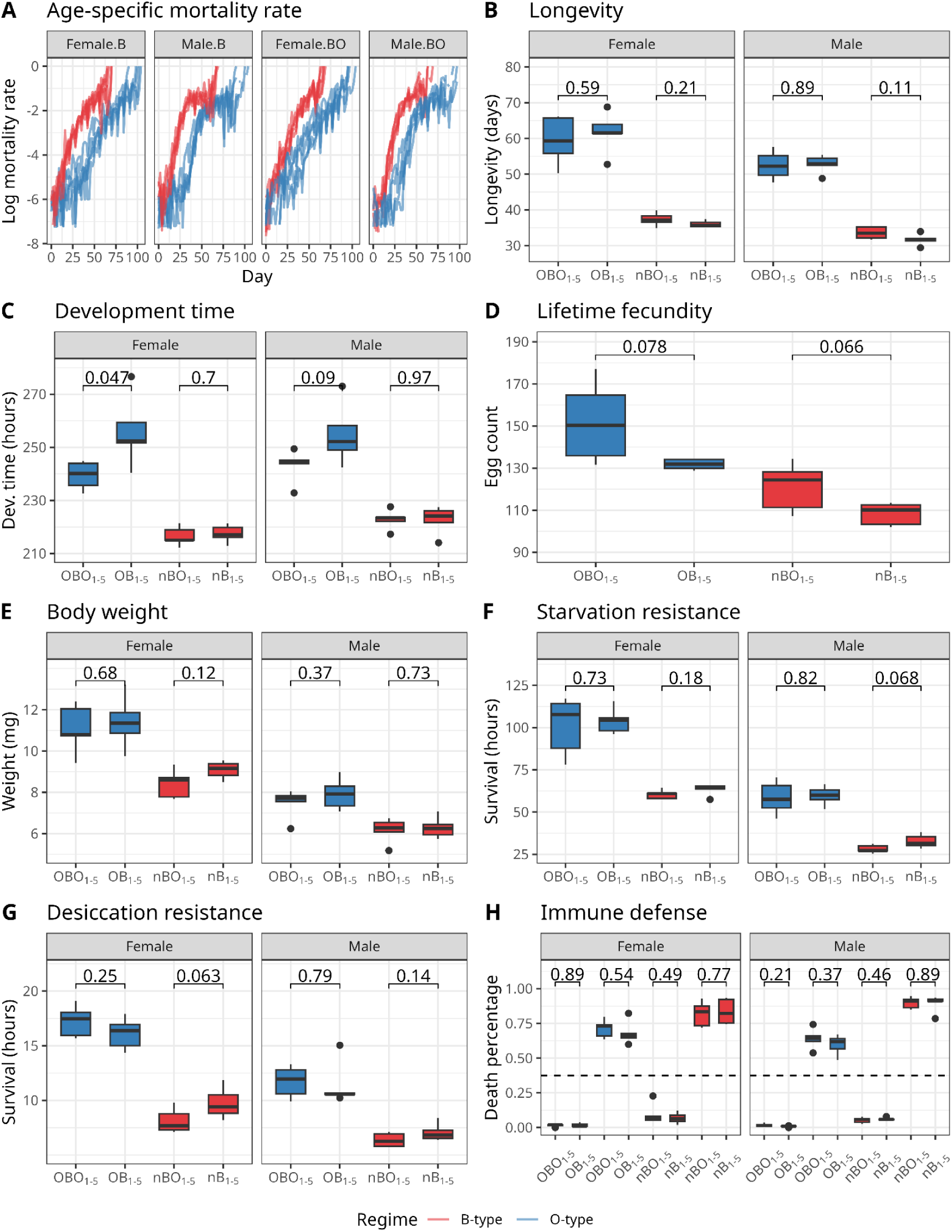
Phenotypic convergence across ancestral backgrounds is observed after 20 generations of O-type selection. Across all panels, data from nBO_1-5_ and nB_1-5_ treatments are shown in red, and data from OBO_1-5_ and OB_1-5_ treatments are shown in blue. Panels B-H, show boxplots derived from 5 data points per group (n=5 for all groups), with each data point representing a sex-specific population mean per replicate population. Male and female data are shown separately where applicable. Statistical differences between ancestries were assessed using t-tests on population means, with p-values reported above comparison brackets. **A** Age-specific mortality rate over time; each line represents one replicate population, with plots split by ancestry (B- or BO-derived) and sex. **B** Mean population longevity (days). **C** Mean development time (hours). **D** Mean lifetime fecundity (average number of eggs laid per female). **E** Mean body weight (mg). **F** Starvation resistance (survival in hours). **G** Desiccation resistance (survival in hours). **H** Immune defense, measured as percent mortality following infection (death percentage); plots above the dashed line show infected files, and plots below the dashed line show uninfected control flies. With the exception of female development time in the OBO vs. OB populations (*p*=0.049) trait values do not significantly differ between ancestral backgrounds. **ALT TEXT:** Graphs comparing phenotype of O-type flies and B-type separated by ancestry, with statistical significance markers.

**Supplementary Figure 3.**
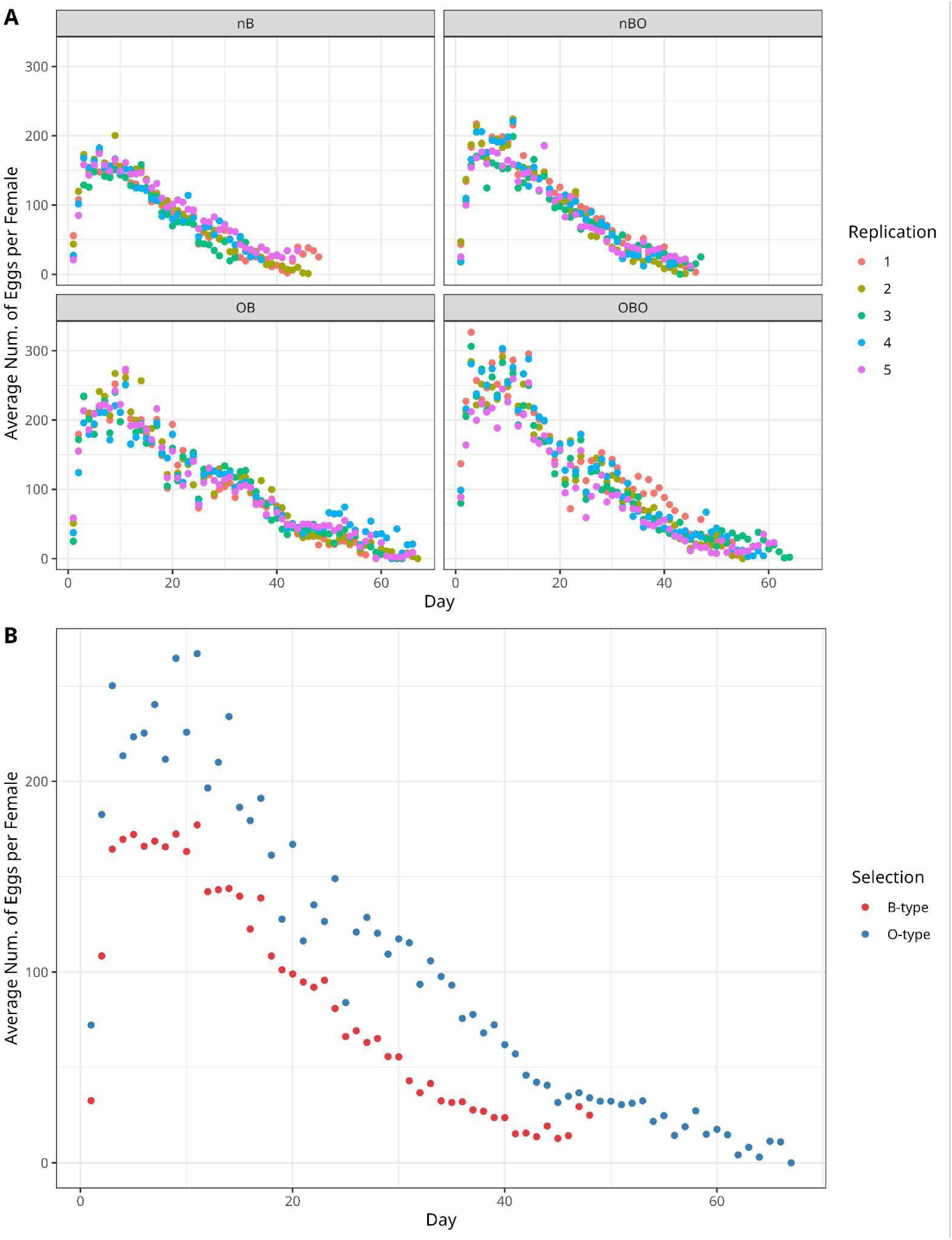
Age specific fecundity trajectories show consistently higher reproductive output in O-type flies across the adult lifespan. Age-specific fecundity was measured at fine temporal resolution after approximately 20 generations of O-type selection. **A** The average number of eggs laid per female is shown per replicate, separated by selection regime (B-type and O-type) and ancestry (B and BO). **B** Age-specific fecundity trajectories averaged across replicates and ancestral backgrounds within each selection regime. **ALT TEXT:** Scatterplots of age-specific fecundity split by ancestry and selection regimen, and split only by selection regimen.

**Supplementary Figure 4.**
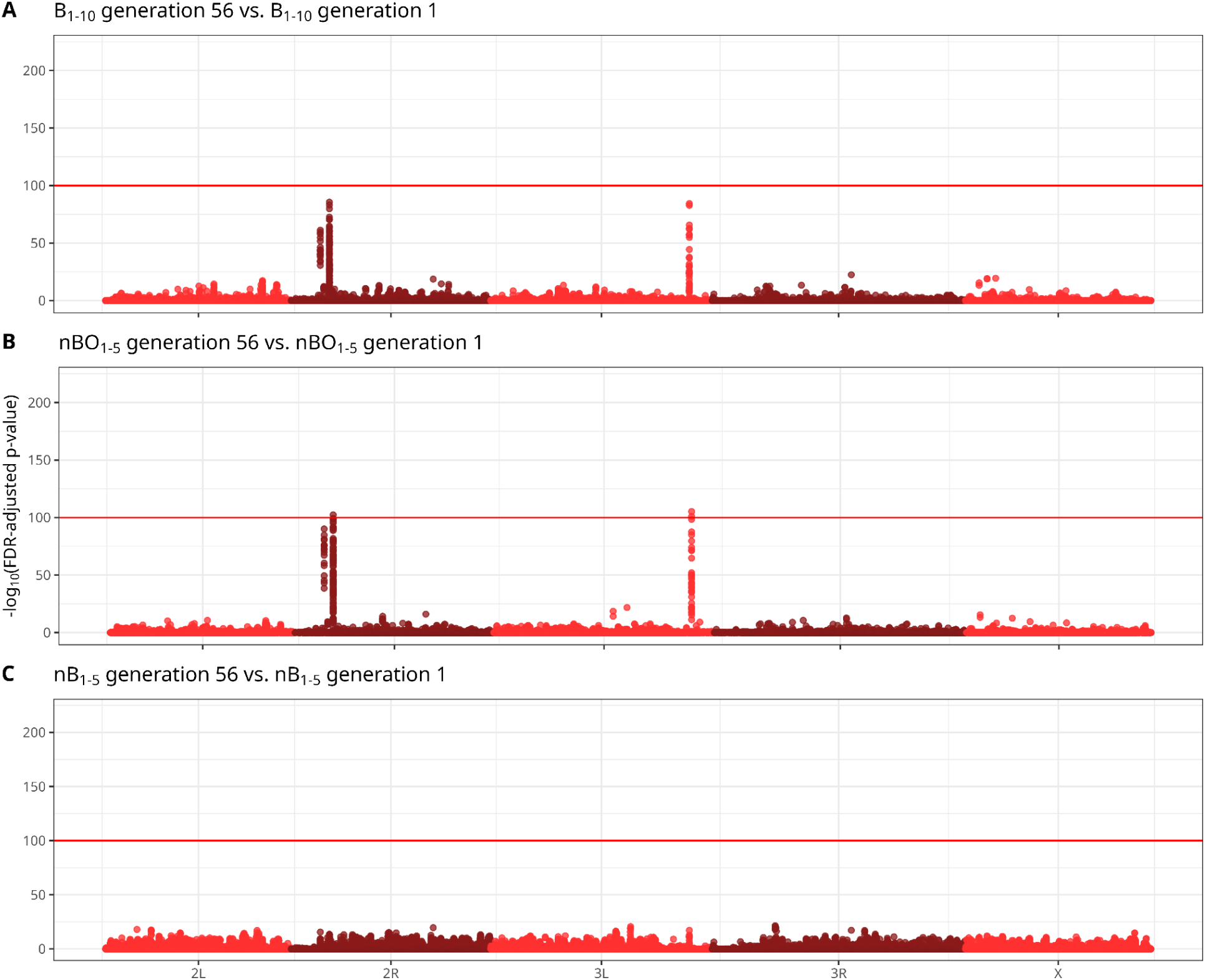
B-type populations show little genome-wide allele frequency change over time, consistent with convergence. Manhattan plots show CMH test results based on scaled SNP frequencies; x-axis values represent cumulative physical distance along the genome and y-axis values represent FDR-corrected *p*-values. The horizontal red line indicates a significance threshold of adjusted *p*-value=10^-100^. **A** The combined ten B-type populations (nBO_1-5_ & nB_1-5_) B_1-10_ at generation 56 vs B_1-10_ at generation 1. **B** nBO_1-5_ at generation 56 vs nBO_1-5_ at generation 1. **C** nB_1-5_ at generation 56 vs nB_1-5_ at generation 1. **ALT TEXT:** Manhattan plots of B-type populations compared between generations 1 and 56, with significance threshold indicating statistically significant single nucleotide polymorphisms.

**Supplementary Figure 5.**
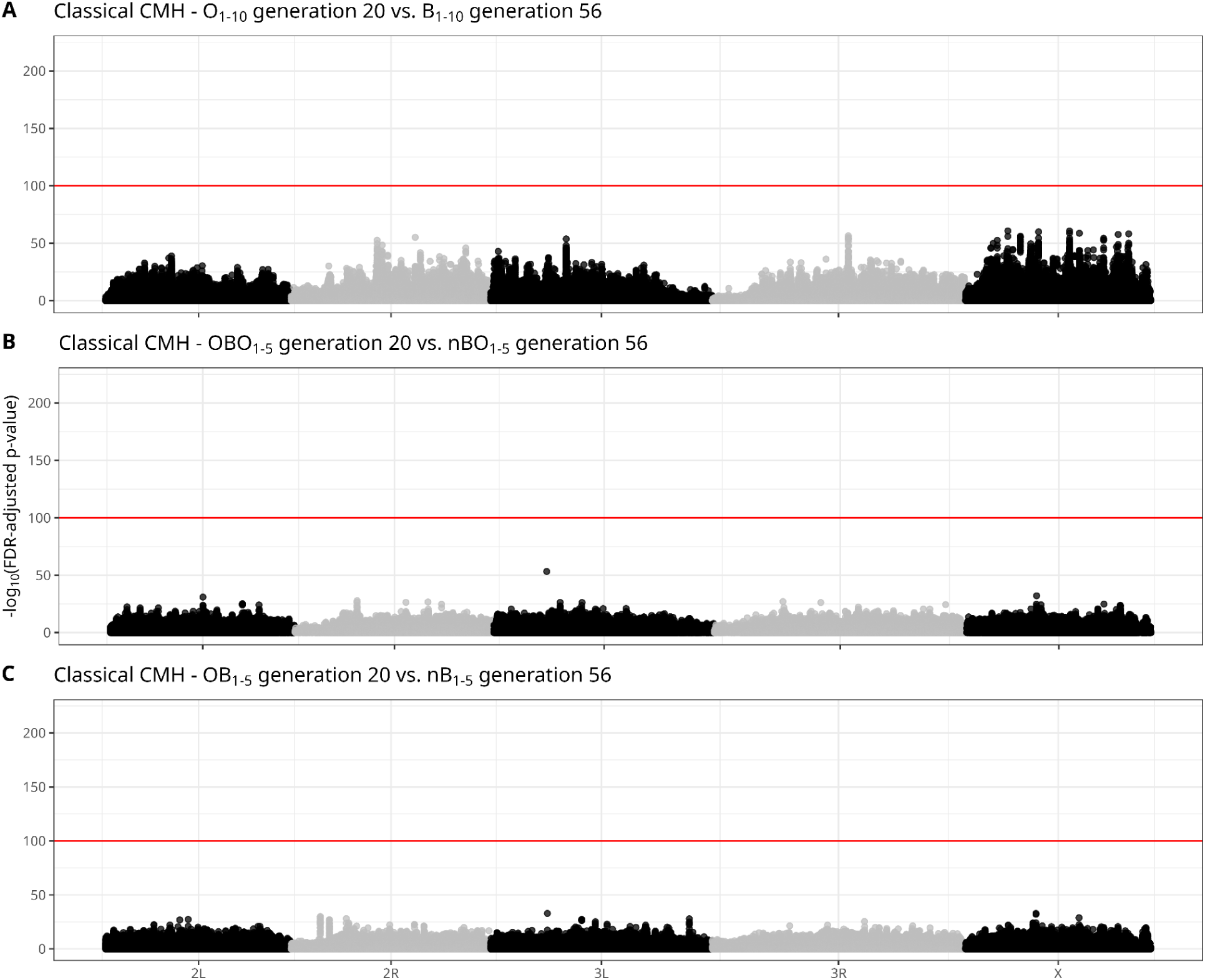
Manhattan plot of adapted CMH tests on scaled allele depth (AD) and unfiltered depth (DP) of all SNPs. SNPs are represented as a function of genomic position and FDR-corrected p-value. The horizontal red line represents a significance threshold of adjusted p-value 10^-100^. **A** The combined ten O-type populations (OBO1-10 & OB1-10) at generation 20 vs the combined ten B-type populations (nBO_1-5_ & nB_1-5_) B_1-10_ at generation 56. **B** OBO_1-5_ at generation 20 vs nBO_1-5_ at generation 56. **C** OB_1-5_ at generation 20 vs nB_1-5_ at generation 56. **ALT TEXT:** Manhattan plots of O-type populations compared to B-type populations, with significance threshold indicating statistically significant single nucleotide polymorphisms.

Supplementary Table 1 - SNPs showing significant allele frequency change under O-type selection and their associated genes

Supplementary Table 2 - Candidate genes underlying the response to O-type selection and their associated GO terms

Supplementary Table 3 - GO terms significantly enriched among candidate genes harboring significant SNPs

Supplementary Table 4 - Statistical model outputs for phenotypic traits analyzed across selection regimes

## References

Ahmad, S. T., Sweeney, S. T., Lee, J.-A., Sweeney, N. T., & Gao, F.-B. (2009). Genetic screen identifies serpin5 as a regulator of the toll pathway and CHMP2B toxicity associated with frontotemporal dementia. Proceedings of the National Academy of Sciences, 106(29), 12168–12173. 10.1073/pnas.0903134106

Araújo, A. R., Reis, M., Rocha, H., Aguiar, B., Morales-Hojas, R., Macedo-Ribeiro, S., Fonseca, N. A., Reboiro-Jato, D., Reboiro-Jato, M., Fdez-Riverola, F., Vieira, C. P., & Vieira, J. (2013). The Drosophila melanogaster methuselah Gene: A Novel Gene with Ancient Functions. PLOS ONE, 8(5), e63747. 10.1371/journal.pone.0063747

Auwera, G. van der, & O’Connor, B. D. (2020). *Genomics in the cloud: Using Docker, GATK, and WDL in Terra* (First edition). O’Reilly Media.

Bagheri, E., Yin, H., Bengo, A. L., Rai, K. E., Conyers, T., Courville, R., Abdoli, M., Burke, M., & Shahrestani, P. (2025). Age-Dependent Immune Defense Against Beauveria bassiana in Long- and Short-Lived Drosophila Populations. MDPI AG. 10.20944/preprints202506.0507.v1

Baldwin-Brown, J. G., Long, A. D., & Thornton, K. R. (2014). The power to detect quantitative trait loci using resequenced, experimentally evolved populations of diploid, sexual organisms. Molecular Biology and Evolution, 31(4), 1040–1055. 10.1093/molbev/msu048

Bioconductor Core Team, B. P. M. O. [Cre. (2017). TxDb.Dmelanogaster.UCSC.dm6.ensGene [Computer software]. Bioconductor. 10.18129/B9.BIOC.TXDB.DMELANOGASTER.UCSC.DM6.ENSGENE

Burke, M. K., Barter, T. T., Cabral, L. G., Kezos, J. N., Phillips, M. A., Rutledge, G. A., Phung, K. H., Chen, R. H., Nguyen, H. D., Mueller, L. D., & Rose, M. R. (2016). Rapid divergence and convergence of life-history in experimentally evolved Drosophila melanogaster. Evolution, 70(9), 2085–2098. 10.1111/evo.13006

Burke, M. K., Dunham, J. P., Shahrestani, P., Thornton, K. R., Rose, M. R., & Long, A. D. (2010). Genome-wide analysis of a long-term evolution experiment with Drosophila. Nature, 467(7315), 587–590. 10.1038/nature09352

Burke, M. K., King, E. G., Shahrestani, P., Rose, M. R., & Long, A. D. (2014). Genome-Wide Association Study of Extreme Longevity in Drosophila melanogaster. Genome Biology and Evolution, 6(1), 1–11. 10.1093/gbe/evt180

Burke, M. K., Liti, G., & Long, A. D. (2014). Standing Genetic Variation Drives Repeatable Experimental Evolution in Outcrossing Populations of Saccharomyces cerevisiae. Molecular Biology and Evolution, 31(12), 3228–3239. 10.1093/molbev/msu256

Burke, M. K., & Rose, M. R. (2009). Experimental evolution with Drosophila. *American Journal of Physiology-Regulatory*, Integrative and Comparative Physiology, 296(6), R1847–R1854. 10.1152/ajpregu.90551.2008

Carlson, M. (2017). *Org.Dm.eg.db* [Computer software]. Bioconductor. 10.18129/B9.BIOC.ORG.DM.EG.DB

Carnes, M. U., Campbell, T., Huang, W., Butler, D. G., Carbone, M. A., Duncan, L. H., Harbajan, S. V., King, E. M., Peterson, K. R., Weitzel, A., Zhou, S., & Mackay, T. F. C. (2015). The Genomic Basis of Postponed Senescence in Drosophila melanogaster. PLOS ONE, 10(9), e0138569. 10.1371/journal.pone.0138569

Chippindale, A. K., Leroi, A. M., Kim, S. B., & Rose, M. R. (1993). Phenotypic plasticity and selection in Drosophila life-history evolution. I. Nutrition and the cost of reproduction. Journal of Evolutionary Biology, 6(2), 171–193. 10.1046/j.1420-9101.1993.6020171.x

Cox, D. R. (1972). Regression Models and Life-Tables. Journal of the Royal Statistical Society: Series B (Methodological), 34(2), 187–202. 10.1111/j.2517-6161.1972.tb00899.x

Di Tommaso, P., Chatzou, M., Floden, E. W., Barja, P. P., Palumbo, E., & Notredame, C. (2017). Nextflow enables reproducible computational workflows. Nature Biotechnology, 35(4), 316–319. 10.1038/nbt.3820

DiAngelo, J. R., Bland, M. L., Bambina, S., Cherry, S., & Birnbaum, M. J. (2009). The immune response attenuates growth and nutrient storage in Drosophila by reducing insulin signaling. Proceedings of the National Academy of Sciences, 106(49), 20853–20858. 10.1073/pnas.0906749106

Djawdan, M., Chippindale, A. K., Rose, M. R., & Bradley, T. J. (1998). Metabolic Reserves and Evolved Stress Resistance in Drosophila melanogaster. Physiological Zoology. (world). 10.1086/515963

Dragojlovic-Munther, M., & Martinez-Agosto, J. A. (2013). Extracellular matrix-modulated Heartless signaling in *Drosophila* blood progenitors regulates their differentiation via a Ras/ETS/FOG pathway and target of rapamycin function. Developmental Biology, 384(2), 313–330. 10.1016/j.ydbio.2013.04.004

Durinck, S., Spellman, P. T., Birney, E., & Huber, W. (2009). Mapping identifiers for the integration of genomic datasets with the R/Bioconductor package biomaRt. Nature Protocols, 4(8), 1184–1191. 10.1038/nprot.2009.97

Fabian, D. K., Garschall, K., Klepsatel, P., Santos-Matos, G., Sucena, É., Kapun, M., Lemaitre, B., Schlötterer, C., Arking, R., & Flatt, T. (2018). Evolution of longevity improves immunity in Drosophila. Evolution Letters, 2(6), 567–579. 10.1002/evl3.89

Foley, E., & O’Farrell, P. H. (2004). Functional Dissection of an Innate Immune Response by a Genome-Wide RNAi Screen. PLOS Biology, 2(8), e203. 10.1371/journal.pbio.0020203

Franceschi, C., Bonafè, M., Valensin, S., Olivieri, F., De Luca, M., Ottaviani, E., & De Benedictis, G. (2000). Inflamm-aging. An evolutionary perspective on immunosenescence. Annals of the New York Academy of Sciences, 908, 244–254. 10.1111/j.1749-6632.2000.tb06651.x

Funakoshi, M., Tsuda, M., Muramatsu, K., Hatsuda, H., Morishita, S., & Aigaki, T. (2011). A gain-of-function screen identifies *wdb* and *lkb1* as lifespan-extending genes in *Drosophila*. Biochemical and Biophysical Research Communications, 405(4), 667–672. 10.1016/j.bbrc.2011.01.090

Graves, J. L., Jr, Hertweck, K. L., Phillips, M. A., Han, M. V., Cabral, L. G., Barter, T. T., Greer, L. F., Burke, M. K., Mueller, L. D., & Rose, M. R. (2017). Genomics of Parallel Experimental Evolution in Drosophila. Molecular Biology and Evolution, 34(4), 831–842. 10.1093/molbev/msw282

Guangchuang Yu. (2018). *Enrichplot* [Computer software]. Bioconductor. 10.18129/B9.BIOC.ENRICHPLOT

Hamilton, W. D. (1966). The moulding of senescence by natural selection. Journal of Theoretical Biology, 12(1), 12–45. 10.1016/0022-5193(66)90184-6

Hervé Pagès, M. C. (2017). *AnnotationDbi* [Computer software]. Bioconductor. 10.18129/B9.BIOC.ANNOTATIONDBI

Hoedjes, K. M., Van Den Heuvel, J., Kapun, M., Keller, L., Flatt, T., & Zwaan, B. J. (2019). Distinct genomic signals of lifespan and life history evolution in response to postponed reproduction and larval diet in Drosophila. Evolution Letters, 3(6), 598–609.

Howell, L., Sampson, C. J., Xavier, M. J., Bolukbasi, E., Heck, M. M. S., & Williams, M. J. (2012). A directed miniscreen for genes involved in the Drosophila anti-parasitoid immune response. Immunogenetics, 64(2), 155–161. 10.1007/s00251-011-0571-3

Irving, P., Troxler, L., Heuer, T. S., Belvin, M., Kopczynski, C., Reichhart, J.-M., Hoffmann, J. A., & Hetru, C. (2001). A genome-wide analysis of immune responses in *Drosophila*. Proceedings of the National Academy of Sciences, 98(26), 15119–15124. (world). 10.1073/pnas.261573998

Ives, P. T. (1970). Further genetic studies of the south amherst population of *Drosophila melanogaster*. Evolution, 24(3), 507–518. 10.1111/j.1558-5646.1970.tb01785.x

Kamakura, M. (2011). Royalactin induces queen differentiation in honeybees. Nature, 473(7348), 478–483. 10.1038/nature10093

Kapahi, P., Zid, B. M., Harper, T., Koslover, D., Sapin, V., & Benzer, S. (2004). Regulation of Lifespan in *Drosophila* by Modulation of Genes in the TOR Signaling Pathway. Current Biology, 14(10), 885–890. 10.1016/j.cub.2004.03.059

Kassambara, A. (2016). ggpubr: “ggplot2” Based Publication Ready Plots (p. 0.6.2) [Dataset]. 10.32614/CRAN.package.ggpubr

Kassambara, A., Kosinski, M., Biecek, P., & Fabian, S. (2025). survminer: Drawing Survival Curves using “ggplot2” (Version 0.5.1) [Computer software]. https://cran.r-project.org/web/packages/survminer/index.html

Kofler, R., & Schlötterer, C. (2014). A Guide for the Design of Evolve and Resequencing Studies. Molecular Biology and Evolution, 31(2), 474–483. 10.1093/molbev/mst221

Lawrence, M., Huber, W., Pagès, H., Aboyoun, P., Carlson, M., Gentleman, R., Morgan, M. T., & Carey, V. J. (2013). Software for Computing and Annotating Genomic Ranges. PLoS Computational Biology, 9(8), e1003118. 10.1371/journal.pcbi.1003118

Libert, S., Chao, Y., Zwiener, J., & Pletcher, S. D. (2008). Realized immune response is enhanced in long-lived *puc* and *chico* mutants but is unaffected by dietary restriction. *Molecular Immunology*, Special Section: Theories and Modelling of T Cell Behaviour, 45(3), 810–817. 10.1016/j.molimm.2007.06.353

Long, A., Liti, G., Luptak, A., & Tenaillon, O. (2015). Elucidating the molecular architecture of adaptation via evolve and resequence experiments. Nature Reviews Genetics, 16(10), Article 10. 10.1038/nrg3937

Luckinbill, L. S., Arking, R., Clare, M. J., Cirocco, W. C., & Buck, S. A. (1984). Selection for delayed senescence in *Drosophila melanogaster*. Evolution; International Journal of Organic Evolution, 38(5), 996–1003. 10.1111/j.1558-5646.1984.tb00369.x

Mandal, L., Martinez-Agosto, J. A., Evans, C. J., Hartenstein, V., & Banerjee, U. (2007). A Hedgehog- and Antennapedia-dependent niche maintains Drosophila haematopoietic precursors. Nature, 446(7133), 320–324. 10.1038/nature05585

McHugh, K. M., & Burke, M. K. (2022). From microbes to mammals: The experimental evolution of aging and longevity across species. Evolution, 76(4), 692–707. 10.1111/evo.14442

Medawar, P. B. (1952). An unsolved problem of Biology.

Obata, F., & Miura, M. (2015). Enhancing S-adenosyl-methionine catabolism extends Drosophila lifespan. Nature Communications, 6(1), 8332. 10.1038/ncomms9332

Partridge, L., & Fowler, K. (1992). DIRECT AND CORRELATED RESPONSES TO SELECTION ON AGE AT REPRODUCTION IN DROSOPHILA MELANOGASTER. Evolution; International Journal of Organic Evolution, 46(1), 76–91. 10.1111/j.1558-5646.1992.tb01986.x

Petersen, A. J., Rimkus, S. A., & Wassarman, D. A. (2012). ATM kinase inhibition in glial cells activates the innate immune response and causes neurodegeneration in Drosophila. Proceedings of the National Academy of Sciences, 109(11), E656–E664. 10.1073/pnas.1110470109

Petrosyan, A., Gonçalves, Ó. F., Hsieh, I.-H., & Saberi, K. (2014). Improved functional abilities of the life-extended Drosophila mutant Methuselah are reversed at old age to below control levels. AGE, 36(1), 213–221. 10.1007/s11357-013-9568-1

Phillips, M. A., Kutch, I. C., Long, A. D., & Burke, M. K. (2020). Increased time sampling in an evolve-and-resequence experiment with outcrossing Saccharomyces cerevisiae reveals multiple paths of adaptive change. Molecular Ecology, 29(24), 4898–4912. 10.1111/mec.15687

Poplin, R., Ruano-Rubio, V., DePristo, M. A., Fennell, T. J., Carneiro, M. O., Auwera, G. A. V. der, Kling, D. E., Gauthier, L. D., Levy-Moonshine, A., Roazen, D., Shakir, K., Thibault, J., Chandran, S., Whelan, C., Lek, M., Gabriel, S., Daly, M. J., Neale, B., MacArthur, D. G., & Banks, E. (2018). Scaling accurate genetic variant discovery to tens of thousands of samples (p. 201178). bioRxiv. 10.1101/201178

R Core Team. (2025). R: A Language and Environment for Statistical Computing. R Foundation for Statistical Computing. https://www.R-project.org/

Remolina, S. C., Chang, P. L., Leips, J., Nuzhdin, S. V., & Hughes, K. A. (2012). GENOMIC BASIS OF AGING AND LIFE-HISTORY EVOLUTION IN DROSOPHILA MELANOGASTER. Evolution, 66(11), 3390–3403. 10.1111/j.1558-5646.2012.01710.x

Rose, M. R. (1984). Laboratory Evolution of Postponed Senescence in Drosophila melanogaster. Evolution, 38(5), 1004–1010. 10.2307/2408434

Rose, M. R., Passananti, H. B., & Matos, M. (2004). Methuselah Flies: A Case Study in the Evolution of Aging. WORLD SCIENTIFIC. 10.1142/5457

Rynes, J., Donohoe, C. D., Frommolt, P., Brodesser, S., Jindra, M., & Uhlirova, M. (2012). Activating Transcription Factor 3 Regulates Immune and Metabolic Homeostasis. Molecular and Cellular Biology, 32(19), 3949–3962. 10.1128/MCB.00429-12

Schlötterer, C., Tobler, R., Kofler, R., & Nolte, V. (2014). Sequencing pools of individuals—Mining genome-wide polymorphism data without big funding. Nature Reviews Genetics, 15(11), 749–763. 10.1038/nrg3803

Schwenke, R. A., Lazzaro, B. P., & Wolfner, M. F. (2016). Reproduction–Immunity Trade-Offs in Insects. Annual Review of Entomology, 61(1), 239–256. 10.1146/annurev-ento-010715-023924

Shahrestani, P., King, E., Ramezan, R., Phillips, M., Riddle, M., Thornburg, M., Greenspan, Z., Estrella, Y., Garcia, K., Chowdhury, P., Malarat, G., Zhu, M., Rottshaefer, S. M., Wraight, S., Griggs, M., Vandenberg, J., Long, A. D., Clark, A. G., & Lazzaro, B. P. (2021). The molecular architecture of Drosophila melanogaster defense against Beauveria bassiana explored through evolve and resequence and quantitative trait locus mapping. G3 *Genes|Genomes|Genetics*, *11*(12), jkab324. 10.1093/g3journal/jkab324

Slack, C., Alic, N., Foley, A., Cabecinha, M., Hoddinott, M. P., & Partridge, L. (2015). The Ras-Erk-ETS-Signaling Pathway Is a Drug Target for Longevity. Cell, 162(1), 72–83. 10.1016/j.cell.2015.06.023

Spitzer, K., Pelizzola, M., & Futschik, A. (2020). Modifying the Chi-square and the CMH test for population genetic inference: Adapting to overdispersion. The Annals of Applied Statistics, 14(1), 202–220. 10.1214/19-AOAS1301

Therneau, T. M., until 2009), T. L. (original S.->R port and R. maintainer, Elizabeth, A., & Cynthia, C. (2024). survival: Survival Analysis (Version 3.8-3) [Computer software]. https://cran.r-project.org/web/packages/survival/index.html

Tong, J. J., Schriner, S. E., McCleary, D., Day, B. J., & Wallace, D. C. (2007). Life extension through neurofibromin mitochondrial regulation and antioxidant therapy for neurofibromatosis-1 in Drosophila melanogaster. Nature Genetics, 39(4), 476–485. 10.1038/ng2004

Troha, K., & Ayres, J. S. (2022). Cooperative defenses during enteropathogenic infection. Current Opinion in Microbiology, 65, 123–130. 10.1016/j.mib.2021.11.003

Tsuzuki, S., Ochiai, M., Matsumoto, H., Kurata, S., Ohnishi, A., & Hayakawa, Y. (2012). Drosophila growth-blocking peptide-like factor mediates acute immune reactions during infectious and non-infectious stress. Scientific Reports, 2(1), 210. 10.1038/srep00210

Turner, T. L., Stewart, A. D., Fields, A. T., Rice, W. R., & Tarone, A. M. (2011). Population-Based Resequencing of Experimentally Evolved Populations Reveals the Genetic Basis of Body Size Variation in Drosophila melanogaster. PLOS Genetics, 7(3), e1001336. 10.1371/journal.pgen.1001336

Vasimuddin, Md., Misra, S., Li, H., & Aluru, S. (2019). Efficient Architecture-Aware Acceleration of BWA-MEM for Multicore Systems. 2019 IEEE International Parallel and Distributed Processing Symposium (IPDPS), 314–324. 10.1109/IPDPS.2019.00041

Walsh, A. K.-G. (2022). *Experimental evolution for longevity differentiation in Drosophila melanogaster*. California State University, Fullerton.

Wang, M. C., Bohmann, D., & Jasper, H. (2003). JNK Signaling Confers Tolerance to Oxidative Stress and Extends Lifespan in *Drosophila*. Developmental Cell, 5(5), 811–816. 10.1016/S1534-5807(03)00323-X

Wang, P.-Y., Neretti, N., Whitaker, R., Hosier, S., Chang, C., Lu, D., Rogina, B., & Helfand, S. L. (2009). Long-lived Indy and calorie restriction interact to extend life span. Proceedings of the National Academy of Sciences, 106(23), 9262–9267. 10.1073/pnas.0904115106

Wickham, H. (with Sievert, C.). (2016). ggplot2: Elegant graphics for data analysis (Second edition). Springer international publishing.

Wickham, H., Averick, M., Bryan, J., Chang, W., McGowan, L., François, R., Grolemund, G., Hayes, A., Henry, L., Hester, J., Kuhn, M., Pedersen, T., Miller, E., Bache, S., Müller, K., Ooms, J., Robinson, D., Seidel, D., Spinu, V., … Yutani, H. (2019). Welcome to the Tidyverse. Journal of Open Source Software, 4(43), 1686. 10.21105/joss.01686

Wickham, H., & Bryan, J. (2015). readxl: Read Excel Files (p. 1.4.5) [Dataset]. 10.32614/CRAN.package.readxl

Williams, G. C. (1957). Pleiotropy, Natural Selection, and the Evolution of Senescence. Evolution, 11(4), 398–411. 10.1111/j.1558-5646.1957.tb02911.x

Yu, G. (2024). Thirteen years of clusterProfiler. The Innovation, 5(6), 100722. 10.1016/j.xinn.2024.100722

Yu, S., Luo, F., Xu, Y., Zhang, Y., & Jin, L. H. (2022). Drosophila Innate Immunity Involves Multiple Signaling Pathways and Coordinated Communication Between Different Tissues. Frontiers in Immunology, 13. 10.3389/fimmu.2022.905370

Zwaan, B., Bijlsma, R., & Hoekstra, R. F. (1995). ARTIFICIAL SELECTION FOR DEVELOPMENTAL TIME IN DROSOPHILA MELANOGASTER IN RELATION TO THE EVOLUTION OF AGING: DIRECT AND CORRELATED RESPONSES. Evolution, 49(4), 635–648. 10.1111/j.1558-5646.1995.tb02300.x.

