## Supplementary Information for "Parallel adaptive responses to postponed reproduction increase lifespan and immune defense"

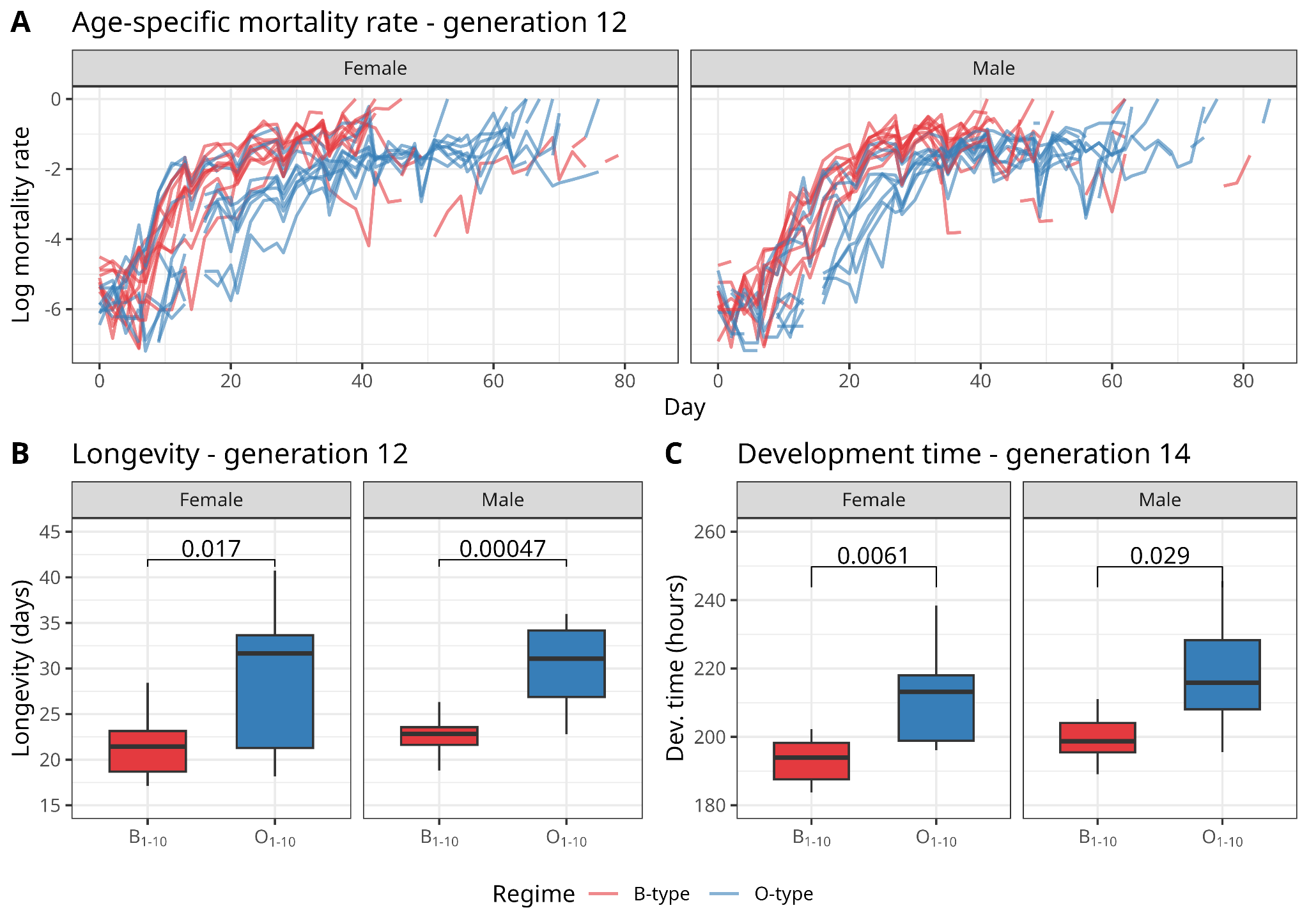

**Supplementary Figure 1. Phenotypic divergence is evident after ~12 generations of O-type selection.** Phenotypic data were collected at an intermediate timepoint, corresponding to generations 12 and 14 of O-type selection and approximately 19 generations of B-type selection. Across all panels, data from the B-type regime (nBO_1-5_ combined with nB_1-5_) are shown in red, and data from the O-type regime (OBO_1-5_ combined with OB_1-5_) are shown in blue. Panels B and C, show boxplots derived from 10 data points (n=10 for all groups), with each data point representing a sex-specific population mean from each replicate population. Data are shown separately for males and females. **A** Instantaneous mortality rate over time; each line represents one replicate population, with plots split by sex. **B** Mean population longevity (days). **C** Mean development time (hours).

**ALT TEXT:** Graphs of phenotypes measured at an intermediate generation time point, with statistical comparisons.

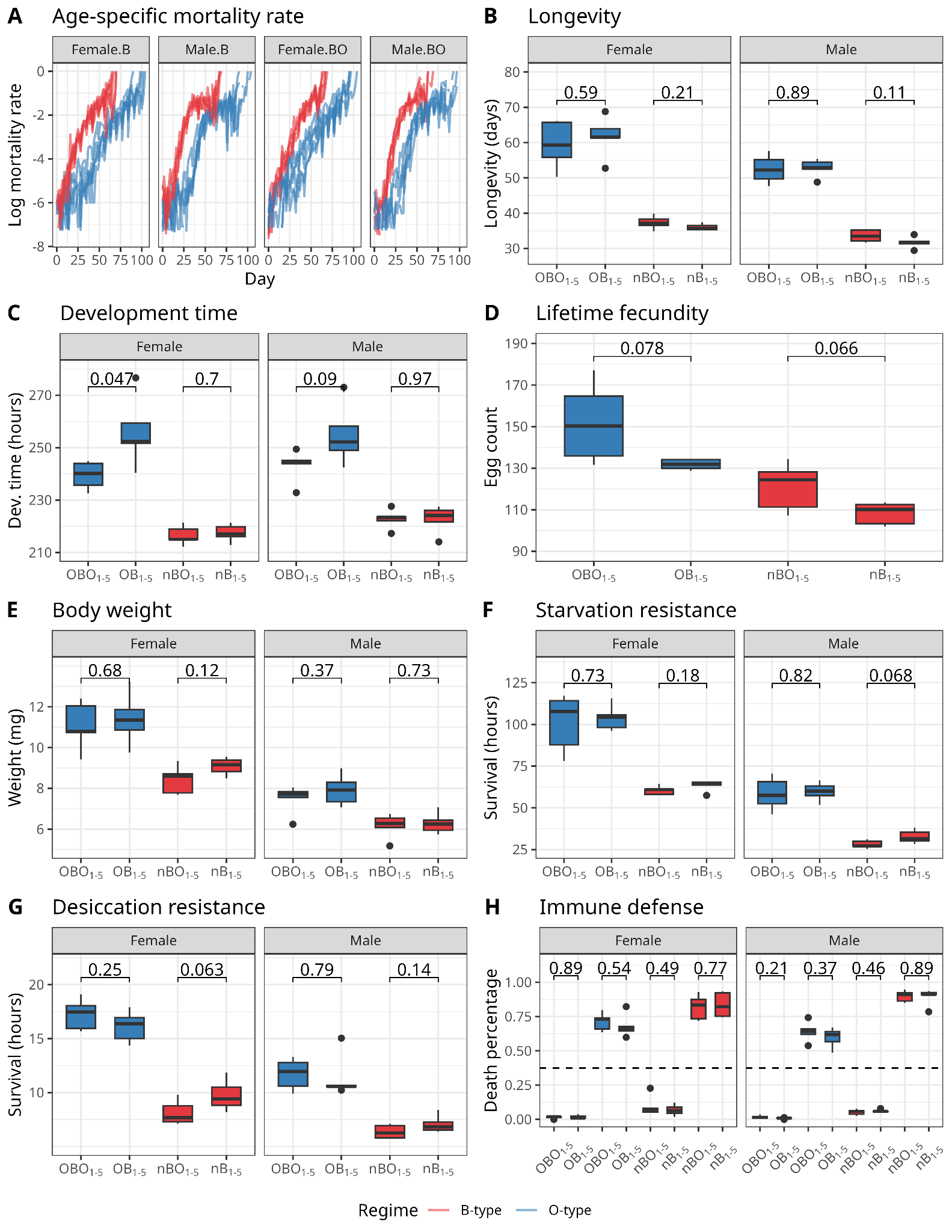

**Supplementary Figure 2. Phenotypic convergence across ancestral backgrounds is observed after 20 generations of O-type selection.** Across all panels, data from nBO_1-5_ and nB_1-5_ treatments are shown in red, and data from OBO_1-5_ and OB_1-5_ treatments are shown in blue. Panels B-H, show boxplots derived from 5 data points per group (n=5 for all groups), with each data point representing a sex-specific population mean per replicate population. Male and female data are shown separately where applicable. Statistical differences between ancestries were assessed using t-tests on population means, with p-values reported above comparison brackets. **A** Age-specific mortality rate over time; each line represents one replicate population, with plots split by ancestry (B- or BO-derived) and sex. **B** Mean population longevity (days). **C** Mean development time (hours). **D** Mean lifetime fecundity (average number of eggs laid per female). **E** Mean body weight (mg). **F** Starvation resistance (survival in hours). **G** Desiccation resistance (survival in hours). **H** Immune defense, measured as percent mortality following infection (death percentage); plots above the dashed line show infected files, and plots below the dashed line show uninfected control flies. With the exception of female development time in the OBO vs. OB populations (*p*=0.049) trait values do not significantly differ between ancestral backgrounds.

**ALT TEXT:** Graphs comparing phenotype of O-type flies and B-type separated by ancestry, with statistical significance markers.

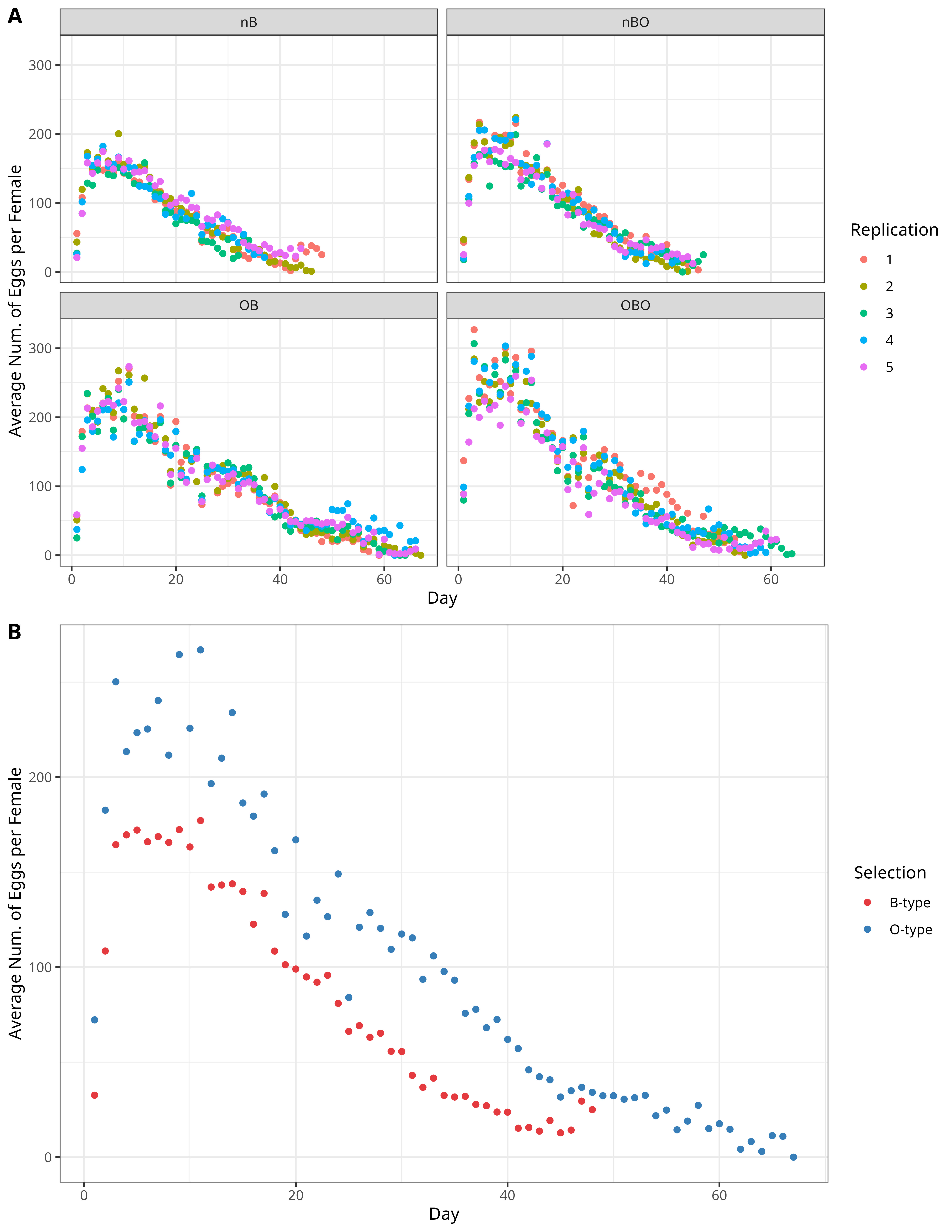

**Supplementary Figure 3. Age specific fecundity trajectories show consistently higher reproductive output in O-type flies across the adult lifespan.** Age-specific fecundity was measured at fine temporal resolution after approximately 20 generations of O-type selection. **A** The average number of eggs laid per female is shown per replicate, separated by selection regime (B-type and O-type) and ancestry (B and BO). **B** Age-specific fecundity trajectories averaged across replicates and ancestral backgrounds within each selection regime.

**ALT TEXT:** Scatterplots of age-specific fecundity split by ancestry and selection regimen, and split only by selection regimen.

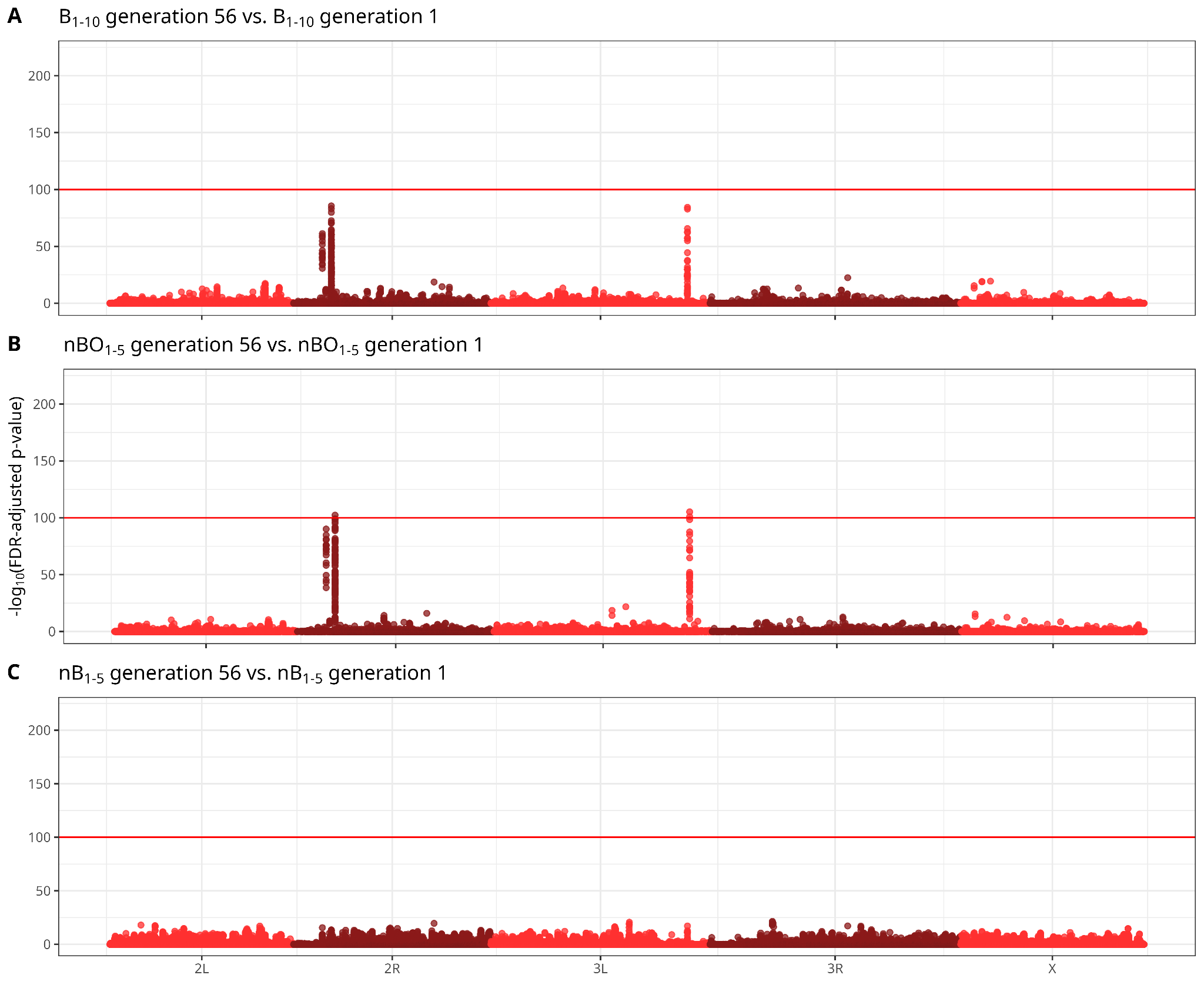

**Supplementary Figure 4.** **B-type populations show little genome-wide allele frequency change over time, consistent with convergence.** Manhattan plots show CMH test results based on scaled SNP frequencies; x-axis values represent cumulative physical distance along the genome and y-axis values represent FDR-corrected *p*-values. The horizontal red line indicates a significance threshold of adjusted *p*-value=10^-100^. **A** The combined ten B-type populations (nBO_1-5_ & nB_1-5_) B_1-10_ at generation 56 vs B_1-10_ at generation 1. **B** nBO_1-5_ at generation 56 vs nBO_1-5_ at generation 1. **C** nB_1-5_ at generation 56 vs nB_1-5_ at generation 1.

**ALT TEXT:** Manhattan plots of B-type populations compared between generations 1 and 56, with significance threshold indicating statistically significant single nucleotide polymorphisms.

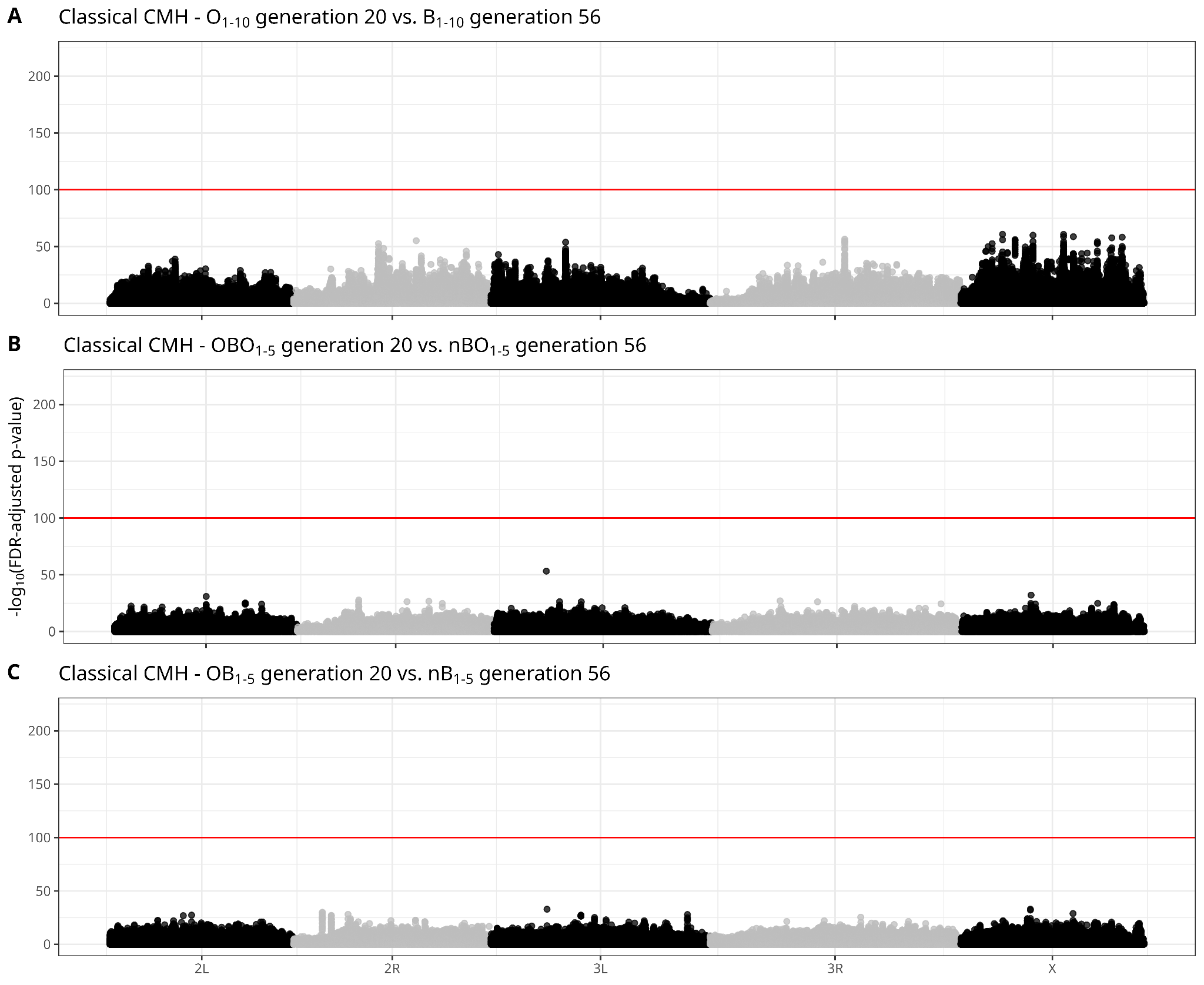

**Supplementary Figure 5.** **Manhattan plot of adapted CMH tests on scaled allele depth (AD) and unfiltered depth (DP) of all SNPs.** SNPs are represented as a function of genomic position and FDR-corrected p-value. The horizontal red line represents a significance threshold of adjusted p-value 10^-100^. **A** The combined ten O-type populations (OBO1-10 & OB1-10) at generation 20 vs the combined ten B-type populations (nBO_1-5_ & nB_1-5_) B_1-10_ at generation 56. **B** OBO_1-5_ at generation 20 vs nBO_1-5_ at generation 56. **C** OB_1-5_ at generation 20 vs nB_1-5_ at generation 56.

**ALT TEXT:** Manhattan plots of O-type populations compared to B-type populations, with significance threshold indicating statistically significant single nucleotide polymorphisms.

Supplementary Table 1 - SNPs showing significant allele frequency change under O-type selection and their associated genes

Supplementary Table 2 - Candidate genes underlying the response to O-type selection and their associated GO terms

Supplementary Table 3 - GO terms significantly enriched among candidate genes harboring significant SNPs

Supplementary Table 4 - Statistical model outputs for phenotypic traits analyzed across selection regimes
